# Homology mediated end joining enables efficient non-viral targeted integration of large DNA templates in primary human T cells

**DOI:** 10.1101/2021.11.12.468427

**Authors:** Beau R. Webber, Matthew J. Johnson, Nicholas J. Slipek, Walker S. Lahr, Anthony P. DeFeo, Joseph G. Skeate, Xiaohong Qiu, Blaine Rathmann, Miechaleen D. Diers, Bryce Wick, Tom Henley, Modassir Choudhry, R. Scott McIvor, Branden S. Moriarity

**Affiliations:** Department of Pediatrics, University of Minnesota, Minneapolis, MN, USA; Masonic Cancer Center, University of Minnesota, Minneapolis, MN, USA; Center for Genome Engineering, University of Minnesota, Minneapolis, MN, USA; Intima Bioscience, New York, USA; Department of Genetics, Cell Biology and Development, University of Minnesota, Minneapolis, MN, USA

## Abstract

Adoptive cellular therapy using genetically engineered immune cells holds tremendous promise for the treatment of advanced cancers. While the number of available receptors targeting tumor specific antigens continues to grow, the current reliance on viral vectors for clinical production of engineered immune cells remains a significant bottleneck limiting translation of promising new therapies. Here, we describe an optimized methodology for efficient CRISPR-Cas9 based, non-viral engineering of primary human T cells that overcomes key limitations of previous approaches. By synergizing temporal optimization of reagent delivery, reagent composition, and integration mechanism, we achieve targeted integration of large DNA cargo at efficiencies nearing those of viral vector platforms with minimal toxicity. CAR-T cells generated using our approach are highly functional and elicit potent anti-tumor cytotoxicity *in vitro* and *in vivo.* Importantly, our method is readily adaptable to cGMP compliant manufacturing and clinical scale-up, offering a near-term alternative to the use of viral vectors for production of genetically engineered T cells for cancer immunotherapy.

## Introduction

Redirecting T cell specificity by introduction of exogenous tumor antigen specific receptor molecules has created a paradigm shift in adoptive cell therapy for cancer. Recombinant DNA technology^1^, the discovery of interleukin-2 (IL-2)^2^, and the development of gammaretroviral vectors converged to allow the first clinical application of stably engineered primary human T cells in the early 1990s^3^. Since then, immense progress has been made both in the identification and cloning of endogenous human T cell receptors (TCR) with inherent tumor antigen specificity, as well as the development of chimeric antigen receptors (CAR)^4^. Currently, most therapeutic cell products are engineered using lenti- or retroviral methods that are not ideal due to high cost, substantial regulatory hurdles, inconsistent viral preps, safety concerns around insertional mutagenesis, and variable transgene expression due to integration site context^5, 6^. In light of these drawbacks, groups have pursued alternative engineering methods. Transient expression of CARs using mRNA has been explored clinically^7^, however the lack of persistent receptor expression may limit ultimate efficacy. Additionally, the use of non-viral DNA transposons, such as *Sleeping Beauty* (*SB*), *PiggyBac* (*PB*), and *TC Buster* (*TCB*), to deliver CARs to T cells for the treatment of cancer has been explored^8–11^. Although DNA transposons improve upon viral methods, there are still concerns regarding insertional mutagenesis and variable transgene expression; moreover, the use of plasmid DNA, required for transposon delivery, is highly toxic to human T cells (<30% viability), necessitating the use of artificial antigen presenting cells (aAPCs) to expand the surviving T cell population in some cases^8, 9^.

Non-viral targeted transgene integration is an appealing alternative, but remains inefficient, especially when integrating larger cargo (>1.5kb)^12–14^. Targeted integration was initially accomplished in mouse embryonic stem cells (mES) using plasmids containing homology to a genomic region of interest for insertion of exogenous DNA sequences through homologous recombination (HR), with integration rates of less than 1×10^-6^ ^15^. The subsequent discovery that genomic double strand breaks (DSBs) can enhance HR by 100 to 1000-fold substantially enhanced the ability to achieve targeted genome engineering^16^. Since this work, several highly active programmable DNA nucleases have been developed for genome engineering, including Zinc finger nucleases, TALENs, and CRISPR/Cas9^17, 18^. DSB induction in primary human cells has largely been optimized by the use of chemically modified gRNAs with the CRISPR/Cas9 system^19^. Use of gRNAs synthesized as RNA oligonucleotides containing 3 tandem 2-O-methyl-3-phosphorothioate modified bases on the 5’ and 3’ ends induced DSBs with efficiencies >90% in T cells and numerous other primary cell types, including hematopoietic stem/progenitor cells (HSPCs)^20–24^. However, targeted insertion of a transgene of interest within these cell populations has been limited by challenges in the delivery of DNA donor template for HR.

Due to the known toxicity of plasmid DNA in primary cells^8, 24, 25^, many researchers have opted to use non-integrating recombinant adeno-associated virus (rAAV) as a DNA donor for HR. The use of rAAV (serotype 6) DNA donor delivery combined with mRNA or protein encoded nucleases has achieved targeting frequencies of >75% in numerous cell types, including HSPC, T, B, and NK cells^21, 24, 26–28^. Although high rates of HR have been achieved in human lymphohematopoietic cells using these methods, there are several substantial drawbacks to this approach. First, the cargo capacity of rAAV (4.5kb) limits the size of expression cassettes that can be integrated^29, 30^. Most DNA donor molecules require homology arms (1kb), a promoter (0.5-1kb), and a polyadenylation sequence (100-250bp), thereby restraining transgene size to 1.75kb-3.4kb. This puts a substantial strain on versatility, especially when integration of multiple genes is required, or when co-integration of a suicide gene is desirable. rAAV approaches to deliver DNA donor molecules also require extremely high amounts of virus, up to 1×10^6^ viral particles per cell, to achieve their unprecedented rates of HR^21, 24, 26–28^. Thus, this approach requires large amounts of expensive, high-quality, high-titer virus, which will be exacerbated when transitioned to cGMP viral production, costing hundreds of thousands of dollars for clinical grade product.

Given that DNA donors typically contain 500-1000bp arms of homology to the desired target site for integration, a universal vector design may be rendered ineffective by the patient’s genetic background. As large numbers of single nucleotide polymorphisms (SNPs) occur in the genome—roughly 1 in every 300bp (ghr.nlm.nih.gov)— there is a strong possibility that SNPs will occur in the targeting arms. Even one SNP close to the transgene cassette in the DNA donor can reduce HR rates by up to 10-fold, and multiple SNPs can reduce this even further^31^. Thus, a one-size-fits-all vector designs will often result in editing rates too low to produce an efficacious therapy if SNPs are present. While it is still possible that this approach may be successful for treating some diseases, such as genetic disorders with a common mutation, there are many other applications that will benefit from patient-specific genome engineering reagents. For instance, one of the most advanced and efficacious methods for cancer immunotherapy employs exomic sequencing of patient tumor samples and identification of cognate tumor reactive TCRs found in the tumor infiltrating T cells^32^. These TCRs are then cloned and delivered to a patient’s T cells using viral delivery, expanded *ex vivo*, and reinfused into the patient to treat their cancer^33^. Alternatively, it would be ideal to integrate the therapeutic TCR transgene as a single copy at a predefined genomic site using HR. While many groups continue to pursue utilizing viral-mediated delivery of donor DNA, the high costs of vector production, limited cargo size, and potential necessity to tailor homology arms to each patient, will certainly maintain this approach as a boutique therapy, only offered at select academic settings, until more cost-effective methods are developed to bring these cutting-edge treatments to all patients in need.

Non-viral genome engineering has been reported in primary human lymphohematopoietic cells using targeted nucleases and DNA plasmid targeting vectors, although the results have been disappointing primarily due to the toxicity of plasmid DNA in these cell types^8, 25^. DNA-induced toxicity has thus fundamentally limited non-viral genome modification in primary cells. We and others^8, 25^ have observed that even high quality, commercially prepared plasmid DNA is toxic to primary human lymphocytes. There have been recent reports using linear PCR derived DNA donor template for efficient (25-50%) non-viral HR in primary immune cells to integrate small cargos (<1kb), however these approaches have suffered from an abrupt loss in efficiency with cargos larger than 1.5kb (<10% efficiency)^12, 14^. It is unclear if this loss in efficiency with larger cargos is due to increased DNA toxicity, inefficiencies of HR, or reduced loading of nucleic acid into cells.

Toxicity of plasmid DNA in primary cells is caused by activation of cytosolic DNA sensors that induce subsequent apoptosis and pyropoptosis^25, 34, 35^. Many cytosolic DNA sensor pathways have been identified and are well characterized^36, 37^. Indeed, previous reports have demonstrated that these pathways and proteins are active and highly expressed in activated human lymphocytes^38^. Some inhibitors have been developed to block these pathways, such as BX795 for TBK1, but the redundancy of these pathways would require the use of numerous inhibitors simultaneously to completely avoid DNA sensing and subsequent cell death^39, 40^. Moreover, many of the proteins appear to be difficult to target using chemical inhibitors, such as cGAS^41^.

Here, we describe methods for efficient CRISPR-based, non-viral engineering of primary human T cells that overcome key limitations of previous approaches, namely DNA-induced toxicity and low efficiency integration of large genetic cargos. By synergizing temporal optimization of reagent delivery, reagent composition, and integration mechanism, we have achieved targeted knock-in of cargo ranging from 1 to 3 kilobases at rates of up to 70% at multiple genomic loci with post-editing cell viability of over 80%; efficiencies nearing those of viral vector platforms. Notably, integration of DNA donor molecules by homology mediated end joining (HMEJ) with short homology arms (48bp) consistently outperformed the use of 1kb homology arms and traditional HR, substantially reducing the potential impact of SNPs on integration efficiencies. As proof of concept, we engineered CAR-T cells and transgenic TCR T cells using a splice acceptor gene construct and gRNA specific to the TRAC locus, such that the CAR or transgenic TCR is expressed under the control of endogenous TRAC regulatory elements. Using this approach, we consistently achieved integration rates of over 35% for CAR-T cells, over 30% for TCR transgenic T cells, and over 25% for super-sized genetic cargo (>6.3Kb). Furthermore, we demonstrate that these cells remain highly functional, maintain low levels of exhaustion markers, excellent proliferation and cytokine production capacity, and demonstrate potent anti-tumor cytotoxicity equal to or better than cells generated using a lentiviral vector. Most importantly, these methods are readily adaptable to cGMP compliant and clinical-scale manufacturing.

In sum, this non-viral genome engineering protocol offers a realistic, near-term alternative to the use of viral vectors in the production of genetically engineered T cells for cancer immunotherapy, providing immense potential for reduced manufacturing time, cost, and complexity without compromising cell expansion or function, while increasing safety and efficacy via targeted integration.

## Materials and Methods

### T Cell Isolation and Culture

PBMCs from de-identified, normal, healthy human donors were obtained by automated leukapheresis (StemExpress). CD4+ and CD8+ cells were sorted by immunomagnetic separation using CD4 and CD8 microbeads (Miltenyi Biotec) in combination with a CliniMACS Prodigy cell sorter (Miltenyi Biotec). Sorted T cells were cultured in OpTmizer CTS medium supplemented with 2.6% Optimize CTS supplement, 5% human AB serum, 1x L-glutamine, 1x Pen/Strep 10mM N-Acetylcysteine, 300U/mL IL2, 5ng/mL IL7, and 5ng/mL IL15 or RPMI, 10%FBS, and 1x Pen/Strep at 37°C and 5% CO_2_ until use. Signed informed consent was obtained from all donors and the study was approved by the University of Minnesota Institutional Review Board (IRB study number 1602E84302). All methods were performed in accordance with the relevant guidelines and regulations.

### Lenti Transduction of T Cells

T cells from 3 donors were thawed and activated using Dynabeads Human T-Activator CD3/CD28 (ThermoFisher) for 24 hours. Post incubation, T cells were treated with Synperonic F 108 (Sigma-Aldrich, 10 µg/mL final concentration) prior to the addition of lentivirus (SignaGen) at a MOI of 20. Cells were then incubated for an additional 24 hours before beads were removed using a magnetic holder and cells were transferred to a G-Rex 24-well plate (Wilson Wolf) in complete OpTmizer CTS medium. Cells were harvested and analyzed for transgene expression 6 days later.

### Non-viral Engineering of T Cells

T cells were thawed and activated using Dynabeads Human T-Activator CD3/CD28 for 24-72 hours. At the specified time, beads were removed using a magnetic holder and washed with PBS once prior to resuspension in P3 electroporation buffer (Lonza). Samples were electroporated using the FI-115 program on the 4D-nucleofector (Lonza) in a 100µL cuvette containing 5-10 × 10^6^ T cells, 5µg sgRNA (IDT), 7.5µg spCas9 mRNA (Trilink), and 5µg of DNA vector (Aldevron) per reaction. Following electroporation, the cuvettes were allowed to rest for 15 minutes before the T cells were gently moved to antibiotic-free medium containing 1µg/mL DNase (StemCell Tech) at 37°C, 5% CO_2_ for 30 min. Following DNase digest, complete CTS OpTmizer T cell Expansion SFM was added and cells were cultured as described above.

### Junction PCR

Primary T cells were integrated with MLSN-CAR or MND-KRAS transgene at the *TRAC1* locus, as well as SA-GFP gene at the AAVS1 locus, through HMEJ and HR mediated CRISPR/Cas9 gene editing. Genomic DNA was isolated from frozen T cell pellets, as well as from wild type controls, using the Thermo Scientific GeneJET Genomic DNA Purification Kit (ThermoFisher Scientific, Waltham, MA). PCR primers were designed targeting both 5’-end and 3’-end of the integration sites, using one primer specific to the insert sequence and the other specific to the genomic sequences outside of the homology arms and HEMJ (**Supplementary Table 1**). PCR was performed using AccuPrimeTaq DNA Polymerase System (ThermoFisher Scientific) with optimized annealing temperature optimized by gradient PCR. Expected amplified fragments were purified using the QIAquick Gel Extraction Kit (Qiagen, Hilden, Germany). Junction PCR products were confirmed by Sanger sequencing (UMGC, U of Minnesota) using PCR primers. Integrants were analyzed by SnapGene version 5.3.0 (SnapGene).

### Polychromatic Flow Cytometry

T cell samples were washed in 1x PBS and incubated with Fixable Viability Dye eFluor780 (eBioscience) for 10 mins at room temperature. Cells were then washed in 1x PBS containing 0.5% BSA and stained with combinations of fluorescently labeled antibodies against CD3, CD4, CD8, CD45ro, CD62L, LAG3, TIM3, PD1, CD25, CD69, Ox40, and 41BB for 15 mins at room temperature (**Supplementary Table 2**). Samples were then analyzed on a CytoFlex S flow cytometer (Beckman Coulter). Data analysis was performed using FlowJo version 10.6.1 (FlowJo LLC).

### Cell Cycle Analysis

T cells were thawed, rested in RPMI1640 supplemented with 10% FBS and 1x Pen/Strep, and then activated using Dynabeads Human T-Activator CD3/CD28. Cells were harvested at 9, 24, 36, 48, and 72 hours, washed and incubated with eFluor780 Fixable Viability Dye eFluor780 (eBioscience) for 10 mins at room temperature before staining with fluorescently labeled antibodies against CD3, CD4, and CD8 for 15 mins at room temperature (**Supplementary Table 2**). Cells were then washed, fixed with -20°C 70% EtOH for 60 mins, and stained with 7AAD (Tonbo Biosciences) and fluorescently labeled antibody against Ki67 (**Supplementary Table 2**) for 15 mins. Samples were then analyzed on a CytoFlex S flow cytometer (Beckman Coulter). Data analysis was performed using FlowJo version 10.6.1 (FlowJo LLC).

### Western Blots

1 × 10^6^ cells were lysed in complete RIPA buffer with protease and phosphatase inhibitors (Sigma-Aldrich). Total protein was quantified using the Pierce BCA Protein Assay Kit (ThermoFisher). 0.5-2 µg/µL of cell lysate was analyzed on the Wes platform (Protein Simple) after denaturation at 95 °C for 5 min. Primary antibodies against actin, STING, pSTING, TBK1, pTBK1, IRF3, pIRF3, IFI16, and AIM2 (Cell Signaling) were used at 1:50-1:100 dilution in kit-supplied buffer and platform-optimized secondary antibodies were purchased from ProteinSimple.

### Coculture and Intracellar Staining

T cells were thawed and cocultured in RPMI1640 supplemented with 10% FBS and 1x Pen/Strep with Raji (CD19+ target cells) or K562 (CD19- target cells) at 1:1 E:T ratio at 37°C with 5% CO_2_ in the presence of a CD107a antibody for 1 hr before treatment with brefeldin A (10 µg/ml; BD Biosciences) and monensin (0.7µg/ml; BD Biosciences). The cells were incubated at 37°C with 5% CO_2_ for 6 hours, harvested, washed twice with 1x PBS and incubated with Fixable Viability Dye eFluor780 (eBioscience) for 10 mins at room temperature. The cells were then washed with 1x PBS and stained with fluorescently labeled antibodies against CD4 and CD8 for 15 mins at room temperature (Supplementary Table X). The cells were then washed again with 1x PBS permeabilized with Fix/Perm (BD Biosciences) for 20 mins at room temperature, washed with Perm/Wash (BD Biosciences) and then stained with fluorescently labeled antibodies against CD3, IFNγ, TNF, and IL2 for 20 mins at room temperature (Supplementary Table X). The cells were then fixed in 1% PFA and analyzed using a CytoFlex S flow cytometer (Beckman Coulter). Data analysis was performed using FlowJo version 10.6.1 (FlowJo LLC).

### In vitro CAR-T Cytotoxicity Assay

T cells were thawed and cocultured in RPMI1640 supplemented with 10% FBS and 1x Pen/Strep with luciferase-expressing Raji target cells at 3:1, 1:1, and 1:3 E:T ratios in triplicate wells at 37°C with 5% CO_2_ for indicted times. Raji cells alone and NP40-treated Raji cells were used as negative and positive controls for targeted killing respectively. At 24 and 48 hours, 5.6 µg of D-luciferin (Goldbio) was added to each sample and bioluminescence was immediately assessed using a BioTek Synergy 2 plate reader running Gen5 software (version 2.04) with an integration time of 1 second. Cytotoxicity was calculated as a percentage of the no-effector control for each E:T ratio.

### Luminex Assay

Cell culture supernatants were collected from the 24-hour time point of the *in vitro* killing assays and clarified by centrifugation to remove possible cell debris (1000xg for 5 minutes). Secreted cytokine analysis was then performed using a ProcartaPlex essential Th1/Th2 human cytokine immunoplex assay according to manufacturer’s instructions (Invitrogen). Immunoassay was then read on a Luminex 200 instrument controlled by BioPlex manager software (version 6.2).

### In vivo CAR-T Therapy Experiments

Specific pathogen-free female NOD-*scid* IL2Rgamma^null^ (NSG) mice were purchased from Jackson Labs. Tumor challenge studies were performed using the Raji-luc cell line, a well-defined model of Burkitt Lymphoma used for testing CAR-T function^42, 43^. Specifically, mice were implanted with 1×10^5^ Raji-luc cells resuspended in 50% Matrigel (Corning) through a 100 µL intraperitoneal (IP) injection. Three days post tumor implantation, the mice were randomized into treatment groups and received either PBS, 5x10^6^ CAR-T cells from 3 donors engineered through viral or non-viral methods as indicated, or 5x10^6^ non-engineered T-cells from matched donors in 100 µL PBS via tail vein injection. Tumor growth was monitored by weekly bioluminescence imaging of mice 5 minutes post IP injection of D-luciferin (100 µL total volume, 28 mg/kg) using an IVIS100 imager followed by ROI analysis of tumor images (Living Image software, version 4.7.3). This study was carried out in strict accordance with the recommendations in the Guide for the Care and Use of Laboratory Animals of the National Institutes of Health. The protocol and all procedures were approved by the University of Minnesota Institutional Animal Care and Use Committee (Protocol 1905-37099A). The health of the mice was monitored daily by University of Minnesota veterinary staff.

### Statistical Analysis

The Student’s t-test was used to evaluate the significance of differences between the two groups. Differences between three or more groups with one data point were evaluated by a one-way ANOVA test. Differences between three or more groups with multiple data points were evaluated by a two-way ANOVA test. Differences between three or more groups with multiple data points were evaluated by Log-rank (Mantel-Cox) test. All assays were repeated in at least three independent donors. Means values ± SD are shown. The levels of significance were set at p<0.05. All statistical analyses were performed using GraphPad Prism 9.2.0.

## Results

### Optimal timing of electroporation post activation is defined by a ‘Goldilocks Zone’

Previous studies have determined that activated primary human T cells are more efficiently transfected by electroporation with nucleic acids compared to resting T cells^44^. In order to determine the optimal timing of this transfection method, we electroporated T cells with mRNA or plasmid DNA encoding EGFP, at 24, 36, 48, or 72 hours post activation with anti-CD3/anti-CD28 Dynabeads and measured GFP expression, proliferation, cytosolic DNA sensor expression, and cell cycle state post vector delivery (**Figure 1A**). Cells were efficiently transfected with EGFP mRNA at all time points (**Figure 1B**). However, cells were more efficiently transfected with plasmid encoding EGFP at 36 hours (47.70% ±12.99) compared to 24 hours (35.56% ±9.08), 48 hours (24.57% ±12.31), and 72 hours (13.35% ±4.86) (**Figure 1B**). This observation correlated with the total number GFP+ cells at each time point 4 days post electroporation; with the 36 hour time point yielding significantly more GFP+ cells than the 24 hour or 72 hour time points (**Figure 1C**).

**Figure 1.**
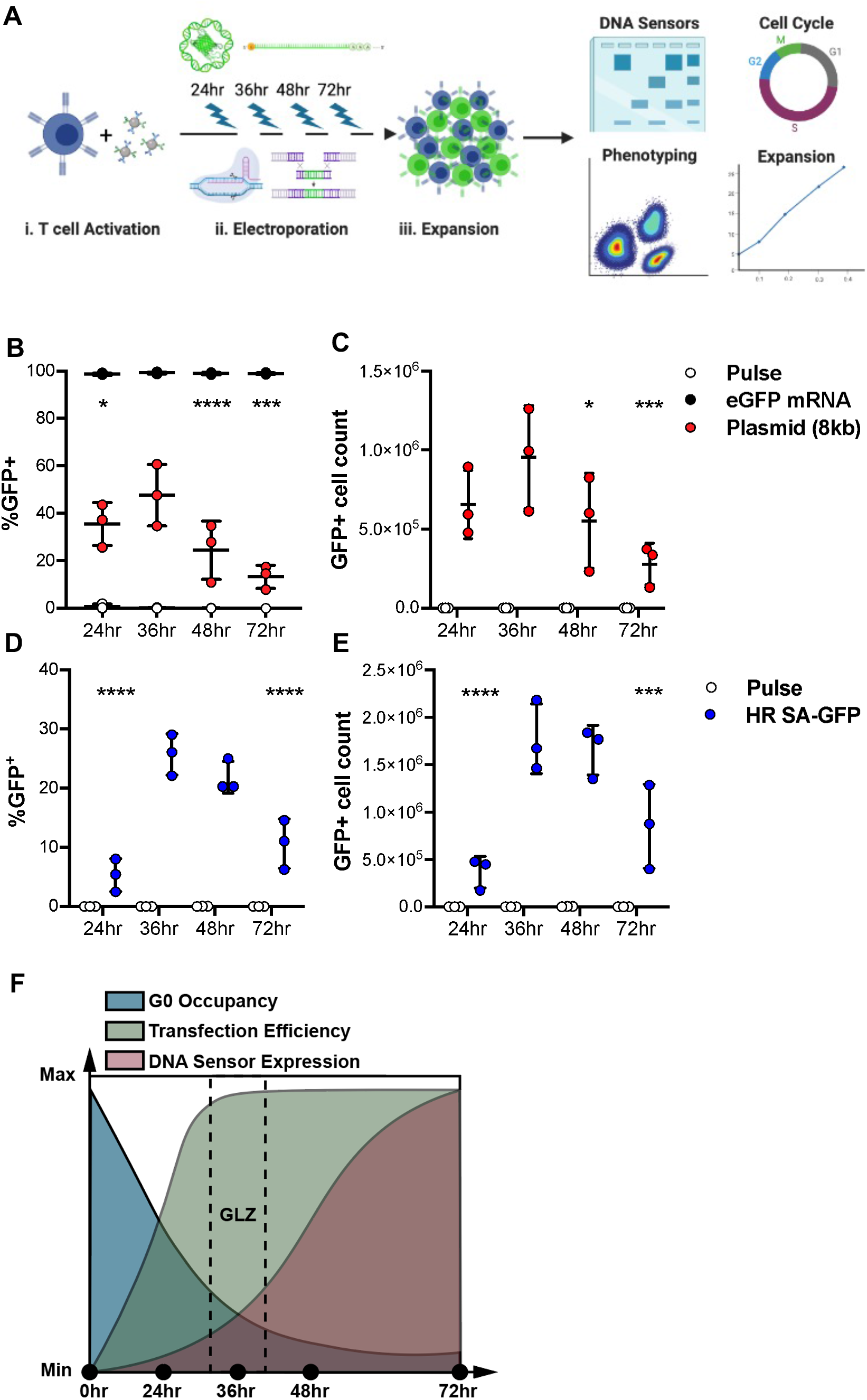
Temporal optimization enhances delivery, expansion, and targeted knock-in in human T cells. **(A)** Diagram of the genome engineering process. ***(i)*** Activation of T cells with anti-CD3/anti-CD28 microbeads. ***(ii)*** Electroporation of cells with plasmid or mRNA encoding GFP, or Cas9, gRNA, and DNA template at one of the indicated timepoints. ***(iii)*** Post engineering expansion of cultures. Percentage of cells expressing GFP **(B)** and total GFP+ cell count **(C)** of cells 4 days after electroporation with mRNA (*black circles*) or plasmid (*red circles*) encoding eGFP compared with pulse-only control (*open circles*) at the indicated time points post activation. Percentage of cells expressing GFP **(D)** and total GFP+ cell count **(E)** 7 days after electroporation with Cas9 mRNA, gRNA targeting AAVS1 and a minicircle plasmid containing a splice acceptor-GFP template with 1000bp homology arms (*blue circles*) compared with no gRNA control (*open circles*) at the indicated time points post activation. **(F)** Diagram of possible temporal factors influencing integration efficiency and cell viability. Unless otherwise noted all statistical analyses were done in comparison to the 36hr timepoint using Two-way ANOVA (n = 3 independent biological donors) (*p<0.05, **p<0.01, ***p<0.001, ****p<0.0001).

Next, we determined the optimal time point to transfect T cells with CRISPR/Cas9-based engineering reagents to stimulate HR by co-delivery of Cas9 mRNA, gRNA targeting the AAVS1 locus, and a minicircle plasmid containing a splice acceptor-GFP template with 1000bp homology arms to the AAVS1 target site. Minicircles are plasmid vectors devoid of the bacterial plasmid DNA backbone and have been shown to improve non-viral engineering of T cells^9, 45^. This construct is designed to express GFP only when successfully integrated into its target location by co-opting endogenous AAVS1 promoter activity. We found that in agreement with our mRNA and plasmid delivery data, cells transfected at 36 hours post activation also had significantly better integration rates (25.7% ±3.5) and total number of edited cells (1.77e6 cells ±3.70e5) than those electroporated at 24 (5.33% ±1.59 and 3.69e5 cells ±1.67e5 respectively) or 72 hours (10.6% ±4.2 and 8.54e5 cells ±4.44, respectively), as assayed by GFP-positivity within populations 7 days post activation (**Figure 1D, E**).

Non-activated T cells in circulation and in culture are in the G0 phase of the cell cycle, a quiescent phase with lowered metabolic activity and gene expression^46^. Moreover, it is also well established that electroporation with nucleic acid, in particular DNA, can be extremely toxic to primary human T cells^8, 25^. This effect is caused, in part, by detection of exogenous cytosolic DNA through cytosolic DNA sensors, which induce apoptosis and pyropoptosis as part of host antiviral defenses^47, 48^. With these points in mind, we first examined the cell cycle states of T cells post-activation using 7AAD and Ki67 staining and also assessed changes in the level and phosphorylation of cytosolic DNA sensors (TBK1, STING, and IRF3) via western blot analysis. Notably, almost all cells were in G0 at the 0 and 24 hour time points post-activation while a majority of T cells had entered the cell cycle at the 36 hour time point and beyond (**Supplementary Figure 1**). This observation coincided with a large increase of the fraction of cells in G1, S, G2, and M phases. Next, we observed that DNA sensors are strongly upregulated at the 48 and 72 hour time points post-activation, which can induce apoptosis and pyropoptosis of cells when activated^47–49^. (**Supplementary Figure 2**). This observation was further confirmed when DNA was included in the electroporation (**Supplementary Figure 2**). Thus, we have identified a critical time point post T cells activation for non-viral engineering, which we call the ‘Goldilocks Zone,’ that maximizes integration due to low-level expression of DNA cytosolic sensors, high frequency of T cells in S/G2 phase of the cell cycle, and high efficiency delivery of nucleic acids (**Figure 1F)**.

### HMEJ allows for high efficiency targeted non-viral integration of large cargo

To further enable non-viral integration of large genetic cargo in T cells, we compared HR and HMEJ approaches using optimal electroporation time points. HMEJ involves targeting a genomic locus with CRISPR/Cas9 and a linearized donor vector with short 48bp homology arms^50^. We and others have demonstrated highly-precise and efficient knock-in using the HMEJ approach in zebrafish, porcine, and human cells^50^. In order to generate the linearized donor template for HMEJ the 48bp homology arms are flanked with two unique CRISPR target sequences, i.e. synthetic universal gRNA sites not found in the human genome. Cas9 binding to these universal gRNA sites results in double-strand breaks that liberate a linearized DNA template from the plasmid. This linearized DNA template is then integrated into the host genome, possibly through DNA repair mechanisms related to Single Strand Annealing (SSA) or Microhomology Mediated End Joining pathways (MMEJ)^51–53^. Here, we present a direct comparison of HMEJ efficacy to the more commonly used HR-based integration strategy (**Figure 2A**).

**Figure 2.**
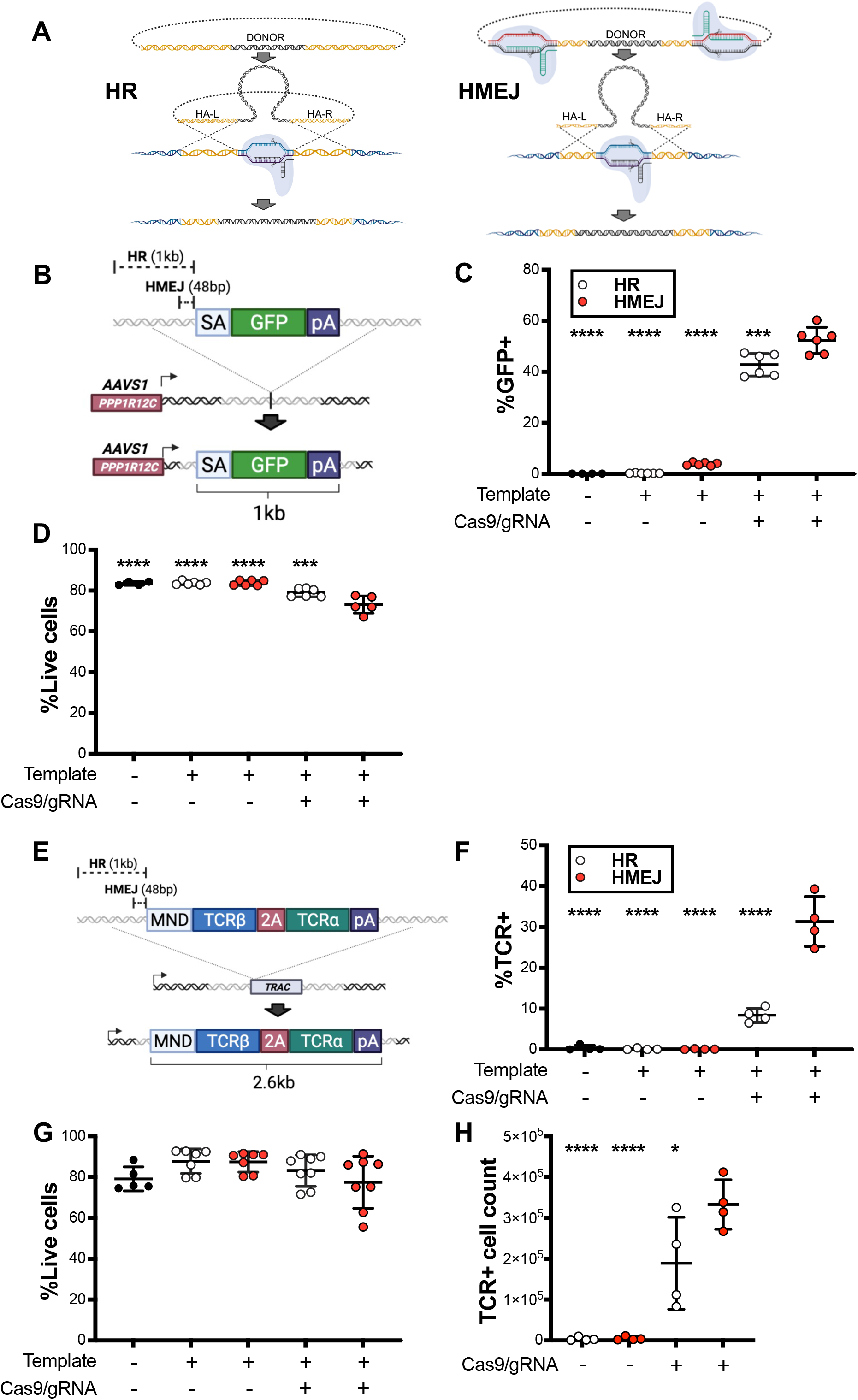
HMEJ mediated integration allows for high efficiency, targeted knock-in. **(A)** Diagrams of HR-mediated (*left panel*) or HEMJ-mediated (*right panel*) DNA integration strategies. **(B)** Diagram of HR and HEMJ templates coding for GFP targeting the *AAVS1* locus. Percentage of cells expressing GFP **(C)** and viability **(D)** 7 days after electroporation with Cas9 mRNA, gRNA targeting either *AAVS1* and minicircle plasmid coding for the GFP under the control of the endogenous promoter using HR (*open circles*) or HMEJ (*red circles*) mediated integration compared with no gRNA and no template controls. **(E)** Diagram of HR and HMEJ templates coding for transgenic TCR targeting the *TRAC* locus. Percentage of cells expressing transgenic TCR **(F)**, viability **(G)** and total transgenic **(H)** TCR expressing cells 7 days after electroporation with Cas9 mRNA, gRNA targeting either *TRAC* and minicircle plasmid coding for the transgenic TCR under the control an MND promoter using HR (*open circles*) or HMEJ (*red circles*) mediated integration compared with no gRNA and no template controls. All statistical analyses were done in comparison to the HMEJ template and Cas9/gRNA samples using One-way ANOVA followed by Dunnett’s multiple comparison test. (n = 5-6 independent biological donors) (*p<0.05, **p<0.01, ***p<0.001, ****p<0.0001).

We designed both HMEJ and HR based templates encoding a splice acceptor-GFP sequence designed to express GFP when correctly integrated under control of the AAVS1 promoter (**Figure 2B**). T cells transfected with HMEJ template, gRNAs and Cas9 were more efficiently engineered (52.4% ±5.16) than those transfected with HR template, gRNA and Cas9 (42.8% ±4.43), but at a cost of a slightly lower viability (79.1% ±2.07 vs 73.1% ±4.25, respectively) (**Figure 2C&D**). We also compared HMEJ mediated integration to another previously reported HR-based integration strategy that uses a linear double stranded DNA template generated through PCR^12, 14^. When using the splice acceptor-GFP construct HMEJ mediated integration resulted in significantly higher GFP expression and total GFP+ cells (32.58% ±0.87 and 9.80e5 cells ±1.31e5, respectively) compared to the linear PCR product approach (9.36% ±2.11 and 6.91e5 cells ±1.31e5, respectively) (**Supplementary Figure 4**).

Next, we tested a much larger, more clinically relevant genetic cargo, the alpha and beta chains of a TCR under control of an MND promoter (**Figure 2E)**, and the difference in integration efficiency between HMEJ and HR was significantly larger (31.35% ±6.12 vs 8.40% ±1.76, respectively) with no significant difference in cell viability (**Figure 2F&G**), and leading to more total TCR+ cells 7 days post activation (3.3e5 cells ±6.1e4 vs 1.9e5 cells ±1.1e5, respectively) (**Figure 2H**). Intriguingly, there were no detectable indels with HMEJ-mediated integration at either the 3’ or 5’ junction, indicating that HMEJ may utilize a more precise form of integration (**Supplementary Figure 3**). These data demonstrate that both HR and HMEJ can mediate efficient integration of small genetic cargo (∼1kb) but efficient integration of large genetic cargo (∼3kb) is much more efficiently achieved using HMEJ.

The two major differences between the HR and HMEJ templates are the size between the homology arm length (1kb vs 48bp) and linearization of the HMEJ vector due to the flanking universal gRNA sites. It is well established that the efficacy of HR mediated integration strongly correlates with the length of homology arms^54^, so we wanted to test the effects of template linearization on integration efficacy. To this end, we developed a ‘Hybrid’ template which has the same 1kb homology arms as the HR template but flanked by the universal target sites from the HMEJ template. We transfected T cells with each of these 3 templates (**Figure 3A, C, E**) along with Cas9 mRNA in the presence or absence of AAVS1 gRNA and the universal target site gRNA. As an additional control we substituted the universal target site gRNA with a truncated 14bp gRNA designed to bind the universal target sites, but not induce a double strand break^12, 55^. T cells transfected with the HR template that included the AAVS1 gRNA demonstrated GFP integration rates of 34.9% ±6.34 (**Figure 3B**). Inclusion of either the universal target site gRNA or the truncated 14bp universal target site gRNA had no significant effect on the integration rate (34.38% ±5.29 vs 38.26% ±12.52, respectively). This result is unsurprising as the HR template lacks the universal target site which allows the universal target site gRNA to bind.

**Figure 3.**
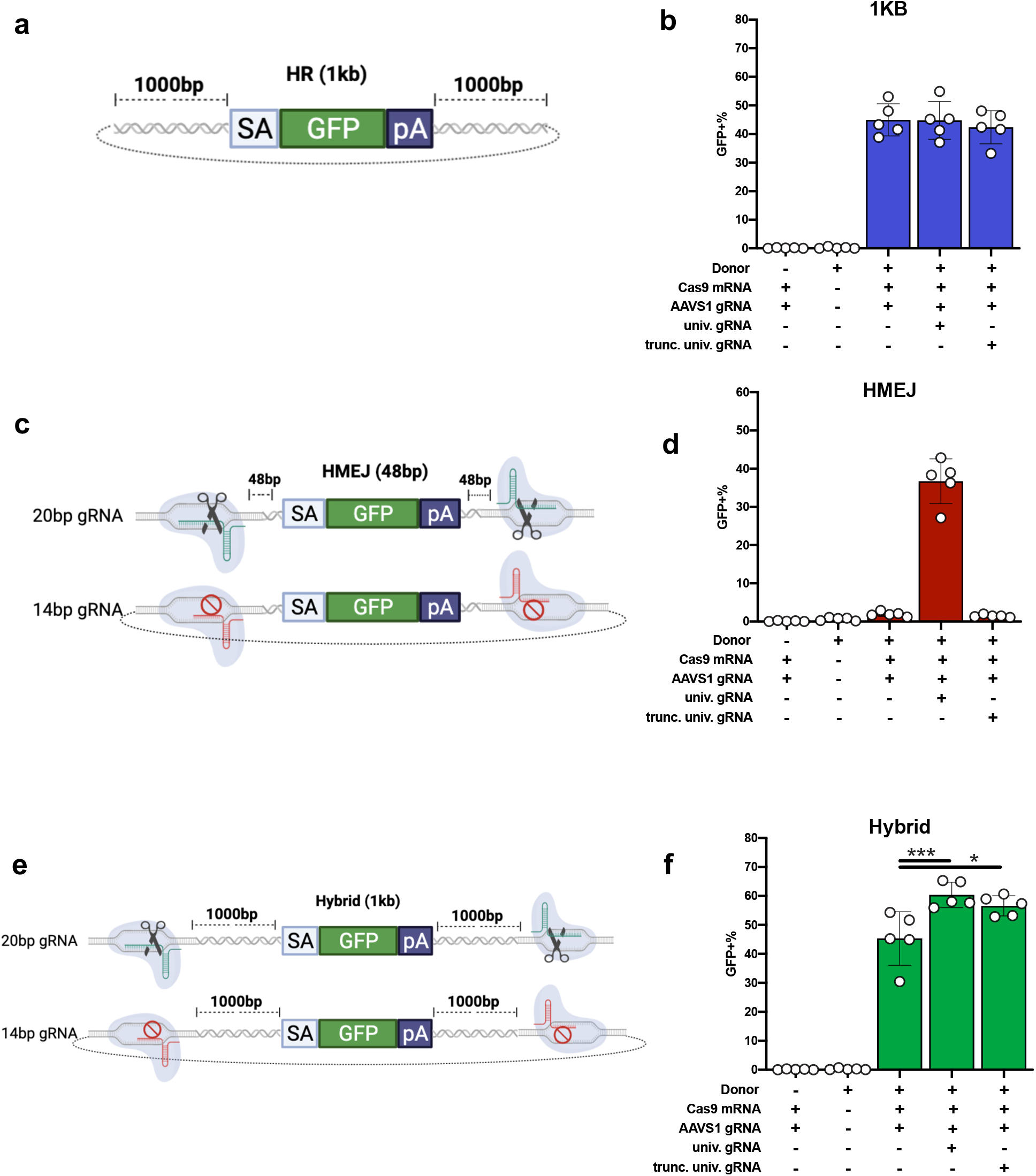
Linearization of template is required for effective integration of short homology HEMJ template. Percentage of cells expressing GFP following electroporation with **(A, B)** 1kb HR template (blue bars), **(C, D)** 48bp HMEJ template (red bars), or **(E, F)** 1kb HMEJ template (green bars) Cas9, AAVS1 gRNA, and either the universal gRNA or a truncated version of the universal gRNA.

T cells transfected with the HMEJ template and AAVS1 gRNA demonstrated low level integration rates (1.71% ±0.67) that were significantly enhanced when the universal gRNA was included (34.13% ±3.77) (**Figure 3D**). Notably, T cells transfected with the truncated 14bp universal gRNA demonstrated low level integration rates (1.46% ±0.34), indicating that Cas9 binding of the template is insufficient to mediate high level integration rates. This demonstrates that linearization of the template with the universal gRNA is likely required for high efficiency HMEJ-mediated integration. In T cells transfected with the hybrid template and AAVS1 gRNA alone we observed integration rates of 45.30% ±9.24 and significantly higher integration in those that also included universal target site gRNA or truncated 14bp universal target site gRNA (60.35% ±4.41 vs 56.57% ±3.47, respectively) (**Figure 3F**). This suggests that binding of the Cas9 protein, which contains nuclear localization signal (NLS) domains, may increase integration by concentrating the associated template in the nucleus.

We also determined the impact of homology arm length with both linearized and non-linearized templates by examining constructs with 48bp, 100bp, 250bp, 500bp, 750bp, or 1000bp flanked with universal target sites in the presence or absence of universal target site gRNAs. While the linearized and non-linearized templates with 1000bp homology arms showed similar levels of integration (62.4% ±6.09 vs 51.5% ±5.84, respectively), non-linearized templates exhibited reduced integration as the homology arms where shortened (9.27% ±1.32 vs 38.75% ±5.29 at 48bp homology) (**Figure 4**). These data demonstrate that HR-based integration rates are significantly diminished when using shorter homology arms, whereas HMEJ-based integration largely maintains integration rates even when using shorter homology arms.

**Figure 4.**
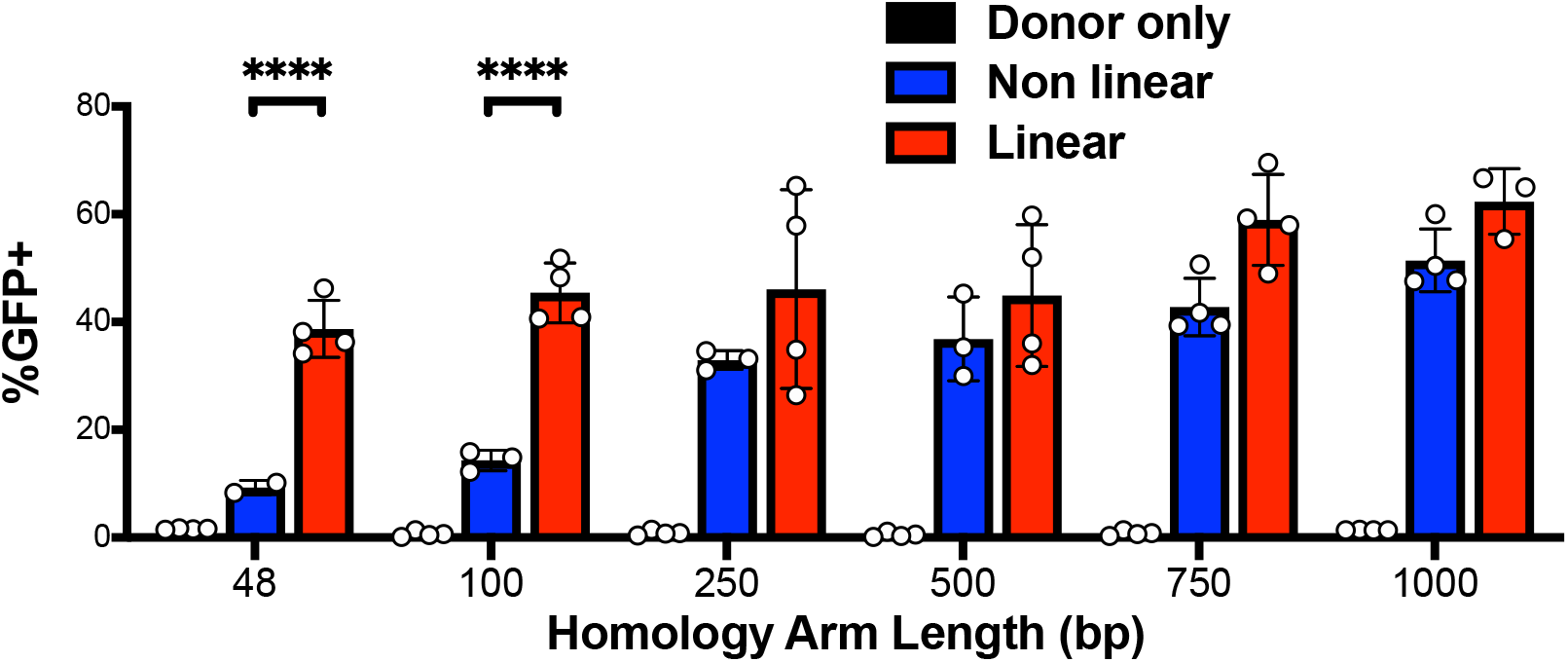
The effect of template linearization on integration increases as homology arms are shorter. Percentage of cells expressing GFP following electroporation with Cas9 mRNA, AAVS1 gRNA, and HEMJ templates with the indicated homology arm lengths. Statistical analyses were done using Two-way ANOVA (n = 3-6 independent biological donors) (*p<0.05, **p<0.01, ***p<0.001, ****p<0.0001).

As HMEJ constructs achieved significantly higher rates of integration with large cargo compared to HR, we wanted to test the ability of HMEJ to integrate super-sized genetic cargo. To this end, we developed a HMEJ-based multicistronic ‘giant construct (GC)’ (6.3kb in size) targeting the TRAC locus. The GC contains an anti-mesothelin CAR and RQR8 sequences under the control of the endogenous TRAC promoter via a T2A element and anti-CD19 CAR, methotrexate-resistant DHFR, and GFP sequences under control of the synthetic MND promoter (**Figure 5A**). RQR8 is a synthetic cell surface molecule containing epitopes derived from CD34 and CD20 that can be used for clinical grade immunomagnetic enrichment (CD34) and targeted suicide using Rituximab (CD20)^56^. T cells were transfected with the GC and CRISPR/Cas9 reagents and assessed for anti-CD19 CAR and GFP expression after 7 days (**Figure 5B**). Remarkably, we were able to achieve >25% integration of the GC using our optimized protocol and HMEJ-based integration (**Figure 5B**). Moreover, methotrexate selection for DHFR expressing T cells further enriched the population of T cells successfully engineered with the GC (**Figure 5B**). Unsurprisingly, the methotrexate selected populations had significantly reduced expansion compared to unselected T cell populations due to eliminations of non-stably integrated T cells in the population. (**Figure 5C**), although the total number of CD19-CAR+ T cells were similar in both groups (**Figure 5D**). Overall, T cells transfected with the GC template and CRISPR/Cas9 reagents expressed anti-CD19 CAR and GFP at 20.1% ±3.43 and 23.3% ±9.79, respectively, and cells selected with methotrexate expressed anti-CD19 CAR and GFP expression at 70.3% ±8.66 and 87.8% ±9.89, respectively. Finally, these GC engineered, methotrexate-selected T cells were also capable of efficiently killing of both mesothelin-expressing A-1847 target cells (**Figure 5E**) and CD19-expressing Raji target cells (**Figure 5F**). Taken together, these data demonstrate that optimized HMEJ-mediated engineering enables efficient integration of super-sized genetic cargo without sacrificing cell viability, proliferation, or functionality.

**Figure 5.**
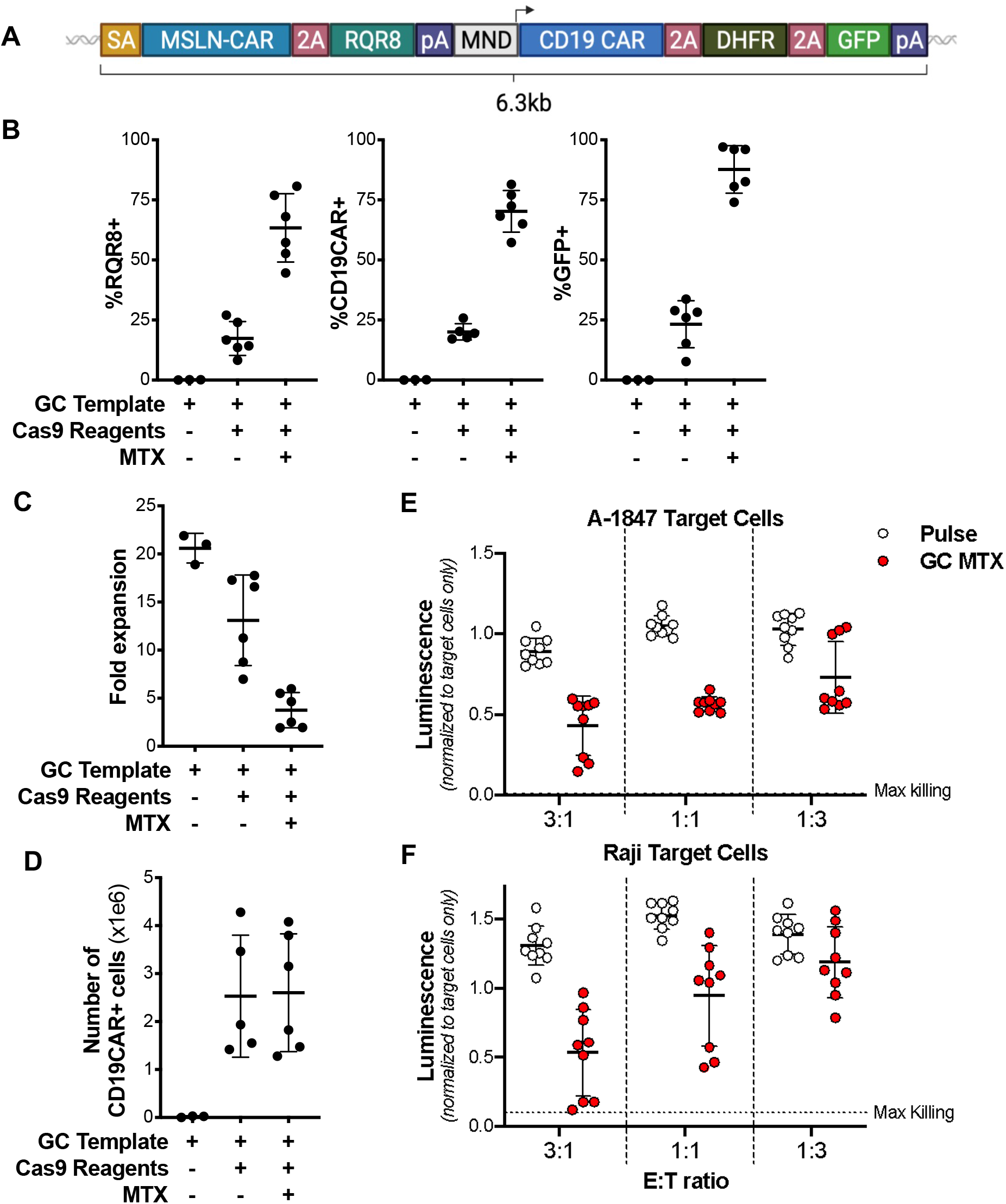
HMEJ allows for efficient integration of very large templates. **(A)** Diagram of the Giant Construct (GC). **(B)** Percentage of cells expressing RQR8 (left panel), CD19-CAR (center panel) or GFP (right panel) 9 days after electroporation with Cas9 mRNA, TRAC gRNA, and GC HMEJ templates in the presence or absence of MTX. **(C)** Fold expansion of cultures and **(D)** total number of CD19-CAR+ T cells after engineering with and without MTX treatment, as well as non-engineered T cells. Luminance of luciferase-labeled A-1847 **(E)** or Raji **(F)** cells follow coculture with GC T cells at the indicated E:T ratio.

### HMEJ engineered CAR-T cells perform comparably to virally engineered CAR-T cells

In order to compare the phenotype and function of HMEJ engineered and virally engineered CAR-T cells we generated CD19 CAR-T cells using HMEJ or a standard lentiviral approach and compared them *in vitro* and *in vivo*. We designed two separate HMEJ constructs encoding a CD19-CAR and RQR8, one containing a T2A element targeting TRAC and the other having a constitutive MND promoter for integration at AAVS1 (**Figure 6A**). As a direct comparison we designed a lentiviral vector with a constitutive MND promoter driving CD19 CAR and RQR8 (**Figure 6A**). T cells from multiple donors were then engineered using our optimized HMEJ approach or industry standard lentiviral transduction (**Supplementary Figure 5)**. Predictably, we observed greater integration rates with lentiviral transduction, as evidenced by RQR8 expression 7 days post engineering (62.9% ±12.7 CD4s, 58.6% ±12.2 CD8s), compared with either non-viral approach (TRAC at 17.0% ±0.85 CD4s, 21.4% ±3.65 CD8s, AAVS1 at 35.3% ±4.66 CD4s, 27.3% ±3.92 CD8s) (**Figure 6B**). Similarly, we observed higher expression, as measured by MFI of RQR8, in the lentiviral transduced cells compared with either the TRAC or AAVS1 engineered cells, likely due to multiple lentiviral integration events (**Supplementary Figure 6**). CD4 and CD8 T cell subsets generated by either non-viral engineering or lentiviral transduction were phenotypically similar, with no statistically significant differences in the percentage of cells in each memory compartment (**Figure 6C**) or deviations in cell surface activation or exhaustion marker expression (**Supplementary Figure 7**).

**Figure 6.**
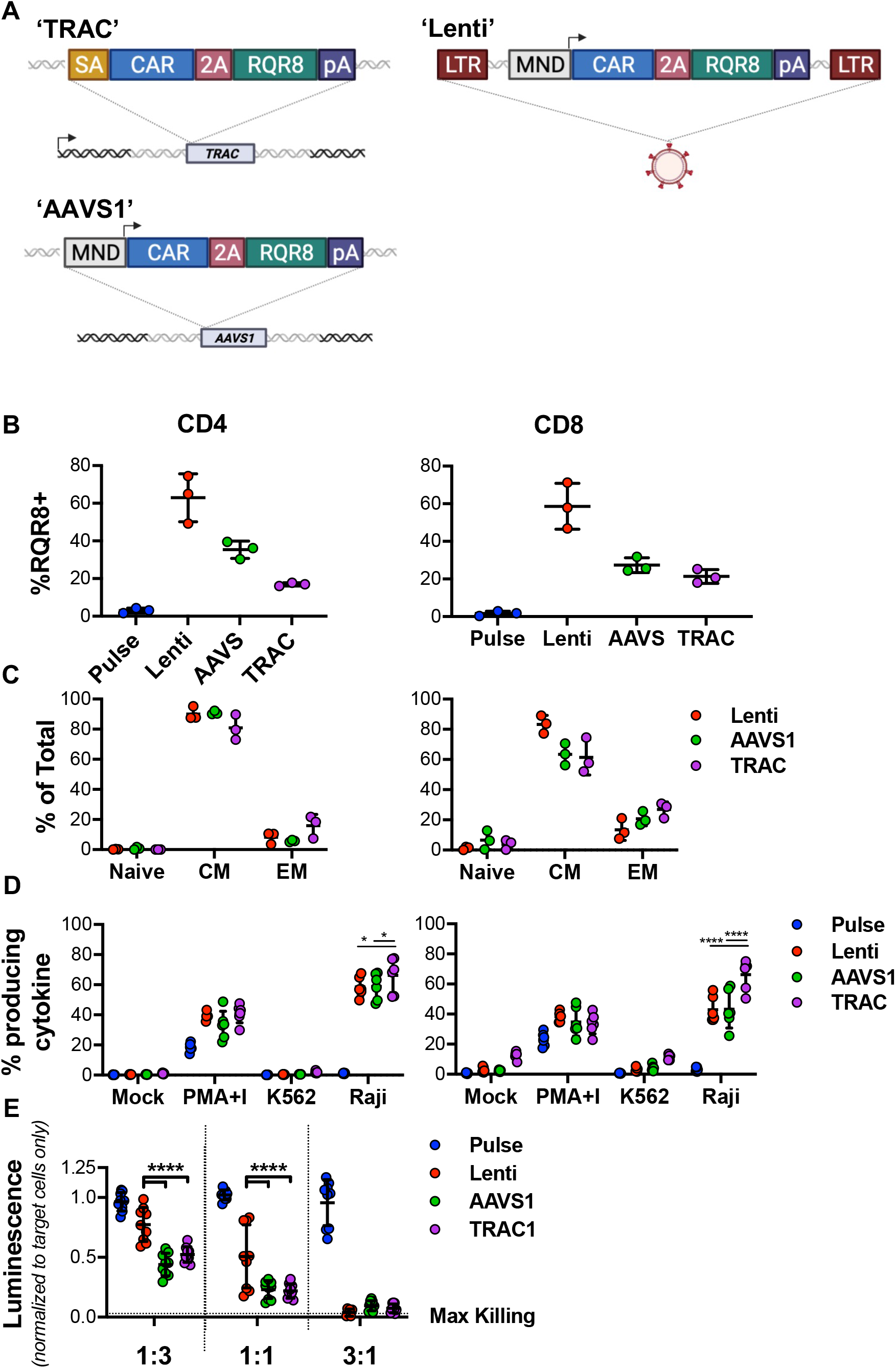
HEMJ engineered CD19 CAR T cells are phenotypically similar to lentiviral transduced cells. **(A)** Diagrams of the constructs used in manufacturing of CD19-CAR T cells. **(A)** Percentage of total T cells expressing RQR8 and **(C)** percentage of cells in naive, central memory, or effector memory compartments after HMEJ non-viral genome engineering or lentiviral transduction with a construct encoding CD19CAR-2A-RQR8. **(D)** Percentage of cells expressing IFNγ, TNF, or IL2 following coculture with K562 cells or Raji target cells, as well as, PMA+Ionomycin treated and no-treatment controls. **(E)** Luminance of luciferase labeled Raji cells follow coculture with CD19CAR T cells engineered with the indicated methods, as well as pulse-control T cells. Statistical analyses were done in comparison to the TRAC engineered cells using One or Two-way ANOVA (n = 3-6 independent biological donors) (*p<0.05, **p<0.01, ***p<0.001, ****p<0.0001).

We next assessed the functionality of the CAR-T cells engineered using non-viral and viral methods by determining their ability to produce cytokines in response to co-culture with CD19 expressing Raji cells (**Supplementary Figure 8**). Through Intracellular Cytokine Staining (ICS) for IFNγ, TNF, and IL2, we found that non-viral TRAC knock-in CAR-T cells had a greater frequency of cytokine expressing cells (IL2+, TNF+, or IFNγ+) (66.22% ±11.47 CD4s, 66.31% ±9.920 CD8s) than those engineered with lentivirus, whereas non-viral AAVS1 knock-in CAR-T cells had similar levels of cytokine production as virally engineered T cells (58.92% ±8.867 CD4s, 43.4% ±12.51 CD8s vs. 58.60% ±6.547 CD4s, 42.94% ±8.554 CD8s) (**Figure 6D**).

Notably, cells transduced with lentivirus produced more TNF, IL4, and IL5 as measured by Luminex assay compared to CAR-T cells engineered with our HMEJ approach at either TRAC or AAVS1, though all three produced similar levels of IFNγ (**Supplementary Figure 9**). As IL4 and IL5 are generally considered T helper 2 (Th2) cytokines this could suggest a bias towards Th2 polarization in lentiviral transduced CAR-T cells when compared to HMEJ engineered CAR-T cells^57^.

We also examined the ability of cells generated with each method to eliminate CD19 expressing Raji cells through a luciferase-based killing assay. After 48 hours of co-culture, cells generated with all three manufacturing methods efficiently killed target cells at the 3:1 ratio. Interestingly, cells manufactured using both the TRAC and AAVS1 HMEJ methods killed a larger percentage of target cells than those manufactured with lentivirus at both the 1:3 and 1:1 effector to target ratios (0.523 ±0.065 and .440 ±0.094 vs 0.775 ±0.141 of max luminescence for 1:3 and 0.219 ±0.060 and 0.230 ±0.075 vs 0.510 ±0.263 of max luminescence for 1:1) (**Figure 6E**). These results demonstrate that CD19 CAR-T cells engineered with non-viral HMEJ approaches are phenotypically and functionally very similar to lentiviral engineered CAR-T cells as assessed by *in vitro* assays.

Finally, given the high efficacy of these CAR-T cells *in vitro* we next examined their ability to clear xenografted tumors *in vivo*. To this end, luciferase expressing Raji cells were used to establish tumors in mice 5-days prior to adoptive transfer of CAR-T cells manufactured using each process. Bioluminescence of tumors was tracked weekly for 56 days to determine tumor growth or remission (**Figure 7A-C**). Importantly, mice receiving CAR-T cells generated with any of the three methods were able to effectively clear established tumors whereas tumor growth continued in untreated mice. All untreated mice bearing Raji tumors had to be euthanized due to tumor burden by day 40 (Median survival of 33.5 days) while only one mouse needed to be sacrificed in the lentiviral engineered CAR-T treated group (**Figure 7D**). Given that this occurred at day 42, well after the mouse had successfully cleared the tumor, we believe this mouse reached one of our defined endpoint criteria due to graft vs host disease (GVHD) rather than continued tumor burden. No other mice showed symptoms of GVHD while under study. Taken together, these data demonstrate that both non-viral HMEJ engineered and lentiviral engineered CAR-T cells are capable of clearing pre-established Raji tumor models in NSG mice.

**Figure 7.**
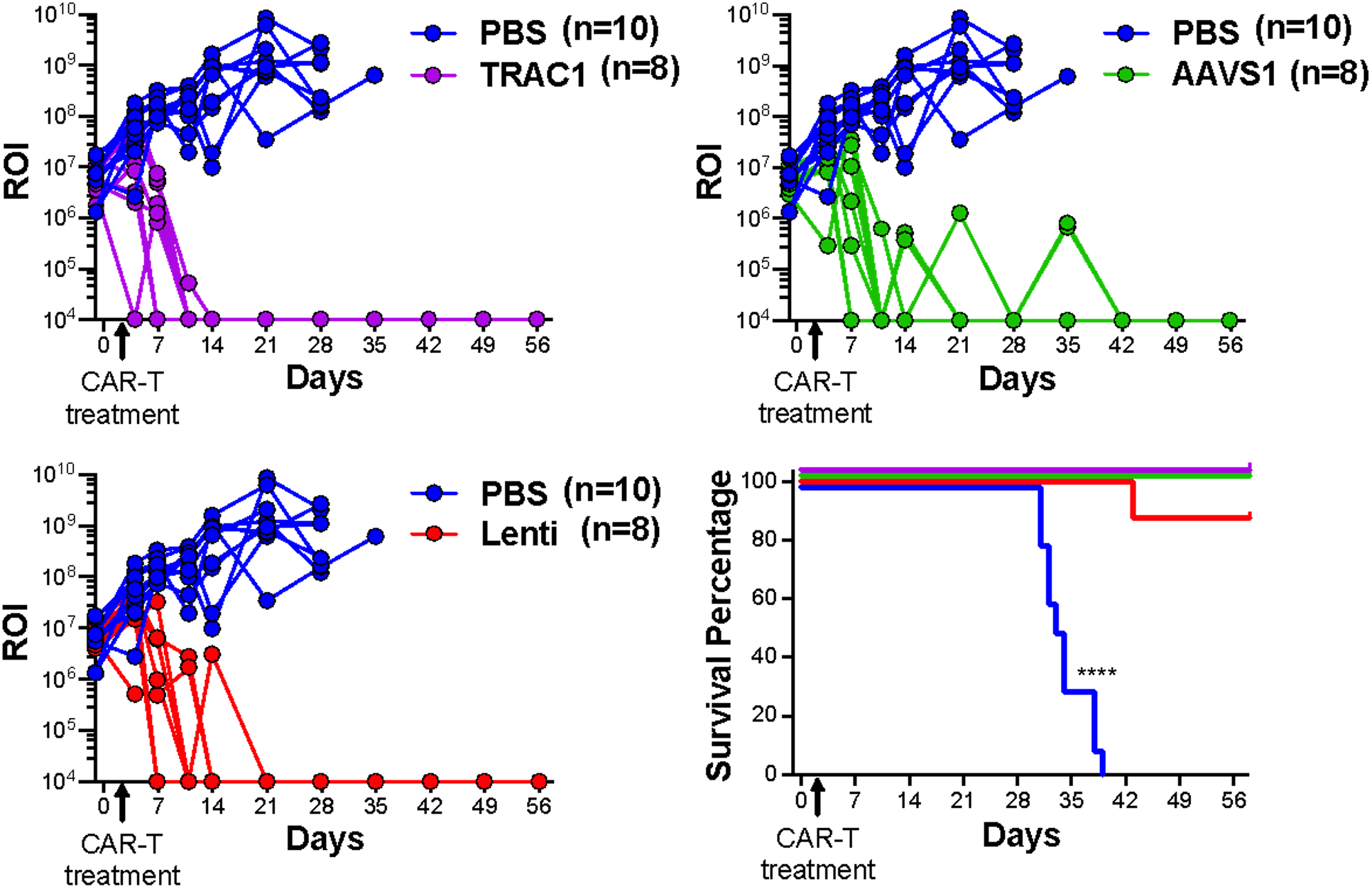
CD19-CAR T cells successfully control Raji tumors *in vivo* regardless of engineering method. (A-C) Luminance (ROI) of individual tumor growth and **(D)** Kaplan Meyer survival of mice bearing Raji tumors following treatment with engineered CD19-CAR T cells compared with mice injected with pulse-only T cells as well as 1x PBS injection-control mice. Log-rank (Mantel-Cox) test was done to compare survival between treatment groups (n = 8-10 animals per group) (****p<0.0001).

## Discussion

As the field of adoptive cell therapy continues to progress, there is a critical need for efficient approaches to engineer immune cells without the cost and complexity of virus-based vectors. In addition, site specific integration is highly desirable as it improves the safety, consistency, and function of engineered cellular products^58^. Although recent studies have successfully demonstrated targeted, non-viral integration of reporter genes and small single CAR/TCR constructs, there is a growing need to use multi-gene expression cassettes to promote enhanced or novel function (e.g. switch receptors^59, 60^, production of cytokines/chemokines and/or associated receptors^59, 61^), improve the safety (e.g. kill switches^62, 63^) and specificity (e.g. logic gate systems^64^) of engineered immune cell products. Thus, it is imperative that non-viral methods are developed for efficient, targeted integration of larger genetic cargo. Here, we identified HMEJ as an efficient mechanism for targeted, non-viral integration in human T cells that is particularly suited for large cargo integration when compared to previously described HR based methods^12–14, 65, 66^. Moreover, we demonstrated that HMEJ can facilitate the integration of large (>6kb), multicistronic (up to 5 genes) expression cassettes at high efficiencies (up to 25%). Such efficiencies were enabled through the discovery that there is a temporal window of time, or Goldilocks Zone, where plasmid-based engineering can take place that minimizes cytosolic-DNA sensor induced toxicity.

In addition to demonstrating large cargo integration, we provide evidence that CD19-CAR T cells engineered using our HMEJ approach are functionally equivalent or better than cells generated with current clinical gold standard lentiviral vector approaches. Specifically, viral and non-viral methods used for CAR-T generation resulted in products that showed similar abilities to kill target cells *in vitro* and produce similar levels of cytokine upon antigen recognition. Notably, and consistent with previous findings^58^, cells generated to express CD19-CAR under the transcriptional control of the TRAC promoter had increased killing and IFNγ production *in vitro*. Importantly though, the cells generated with either method were equally able to clear established CD19-expressing tumors in an *in vivo* NSG model.

This work establishes that there is a critical window post activation where T cells are highly amenable to transfection, but do not experience the severe cytosolic DNA-induced cytotoxicity seen at later time points following activation. These factors combine to make the 36-hour time point ideal for non-viral genome engineering of primary human T cells, which we have coined as the ‘Goldilocks Zone.’ Circulating peripheral immune cells, particularly T and B cells, can be remarkably quiescent, with very little metabolic activity, a factor that may contribute to their low transfection efficiency pre-activation^67^. Many other primary cell types, such as NK cells and B cells, require activation for long term culture and expansion^21, 28^. It is unclear if a similar Goldilocks Zone for non-viral genome engineering might exist for these cells as well.

Another appreciable finding of the current study is that our HMEJ genome engineering approach performs significantly better than HR-based approaches with larger genetic cargo. This has major implications on eventual use in the development of cellular products for clinical use as the majority of therapeutic constructs such as 2nd generation CARs (∼1.6Kb) and TCRs (∼2Kb) are considerably larger than the reporters used in many studies, such as GFP (∼700bp) or RQR8 (∼500bp). It is uncertain why HMEJ allows for greater integration than HR at larger cargo sizes, but there are several possible contributing factors. The 48bp homology arms required for HMEJ mediated integration are much shorter than the more standard 1Kb homology arms required for efficient HR mediated integration. This allows for the removal of almost 2Kb of sequence in the plasmid vector, greatly reducing the length and mass of the delivery vector. Thus, the same mass of the HMEJ construct contains many more copies of the template minicircle than the HR construct, therefore, the stoichiometric ratio of template copies per cell is much higher for HMEJ than HR. It is not possible to simply increase the mass of HR template used in manufacturing in order to equalize the template copy numbers as increasing the amount of template DNA further increases DNA-induced cell apoptosis^68^. Another possible factor impacting integration rates could be the linear template generated during the HMEJ integration strategy. This may allow the template to better mimic the linear sister chromatid normally used during repairs of double stranded DNA breaks. Furthermore, it has been previously shown that including truncated gRNA binding sites on the DNA donor template allow Cas9 to bind and drag it into the nucleus via the NLSs^12^. This combination of linearized template with increased nuclear localization of the donor template may thus synergize, resulting in higher rates of large cargo integration with HMEJ.

The mechanism of HMEJ-meditated DNA integration is not currently well defined. Previous work has suggested that it is more likely mediated by alternative-end joining/microhomology-mediated end joining (MMEJ), or by single strand annealing (SSA) DNA repair pathways as opposed to nonhomologous end joining (NHEJ)^50, 69^. Strikingly, we were unable to detect any indels at either the 3’ or 5’ integration sites, indicating the error prone NHEJ pathway is likely not involved with HMEJ. Moreover, the lack of indels and consistency of the integration event is ideal for downstream clinical applications.

A major advantage of using a non-viral approach to genetically engineer T cells for high-quality, cGMP-compliant cell therapy products is the potential for greatly reducing the time, cost, and complexity of manufacturing lentivirus. Preparing the clinical grade lentiviral vector alone is very expensive and can take up to 18 months to produce and quality check^70, 71^. This is prohibitively expensive and time-consuming for widespread use in patient-specific constructs for cellular immunotherapies. In contrast, the generation of cGMP-quality minicircle plasmid template for use in our non-viral approach allows for rapid development and deployment of cellular products to the bedside on a much faster and efficient timeline^72^.

Current clinical studies using a non-viral approach to engineer CAR-T cells largely utilize DNA transposon systems to introduce the CAR construct into T cells (Clinical trial numbers NCT03389035, NCT04289220). While the use of transposons can allow for high level integration rates, on par with lentivirus, their use is limited by their mechanism of integrating at random locations within the genome^73–77^. This causes significant drawbacks, including the potential for integration into tumor suppressor genes or oncogenes, imprecise control of the number of copies integrated per cell, and an inability to use endogenous promoters and their regulatory elements to control expression of the integrated construct. In contrast, CRISPR/Cas9-based genome engineering allows for precision insertion of the construct, both allowing for control of the copy number and the ability to use endogenous promoters to control expression. This specific integration and lack of detectable off-target editing reduces or eliminates concerns of adverse events due to mutations within tumor suppressor genes or oncogenes. Furthermore, precision engineering allows for simultaneous knock-in/knock-out engineering strategies where the construct is inserted into and disrupts a target gene, thus achieving the knock-in and knock-out simultaneously. Moreover, knocking out additional genes in parallel simply requires additional gRNAs targeting those genes to be included in the electroporation mix, though this approach may lead to increased risk of chromosomal aberrations and translocations^20^.

Importantly, our HMEJ method is readily compatible with cGMP compliant manufacturing and clinical scale-up. The methods and reagents used for isolating, culturing, activation, and expanding T cells are routinely used in clinical trials and clinical products^78–80^. For instance, the Cas9 mRNA is cGMP compliant and already used in clinical trials (NCT04426669), both the ugRNA and our targeting gRNA are available in cGMP compliant forms and are shown to induce minimal to no off target editing^50, 58^. Additionally, the methods for generating minicircles has already been adapted to generate cGMP-quality reagents at clinical scales, without the expense or difficulty required for other DNA template preparations^72^.

Furthermore, a major advantage of our HMEJ manufacturing over other published non-viral engineering approaches is the ability to achieve high-level integration of super-sized cargo while maintaining high cell viability and functionality^12–14, 65, 66^. Alternative approaches have relied on hard-to-produce linear DNA or ssDNA templates and are limited to smaller cargos of low sequence complexity^14^. These limited small cargo-size approaches allowed for highly efficient integration of promoterless 2nd generation CAR constructs (∼1.6Kb), but appear to struggle with larger TCR constructs or strategies involving the integration of multiple proteins (e.g. multiple CARs, CAR+selection marker, or CAR+modified cytokine/chemokine receptor)^12–14, 65, 66^.

Here, we demonstrate a method, based on HMEJ, for generating CAR/TCR-T cells that remain highly functional while retaining low expression of exhaustion markers, excellent proliferation capacity, and potent anti-tumor cytotoxicity equal to or better than cells generated using viral vectors. Most importantly, these methods are readily adaptable cGMP compliant and clinical scale-manufacturing even with super-sized cargo. This genome engineering method offers a realistic, near-term alternative to the use of viral vectors for production of genetically engineered T cells for cancer immunotherapy. It holds great potential for reduced manufacturing time, cost, and complexity in comparison to viral vectors while increasing safety and efficacy through their site-specific nature.

## Supplementary Figure Legends

**Supplementary Figure 1.**
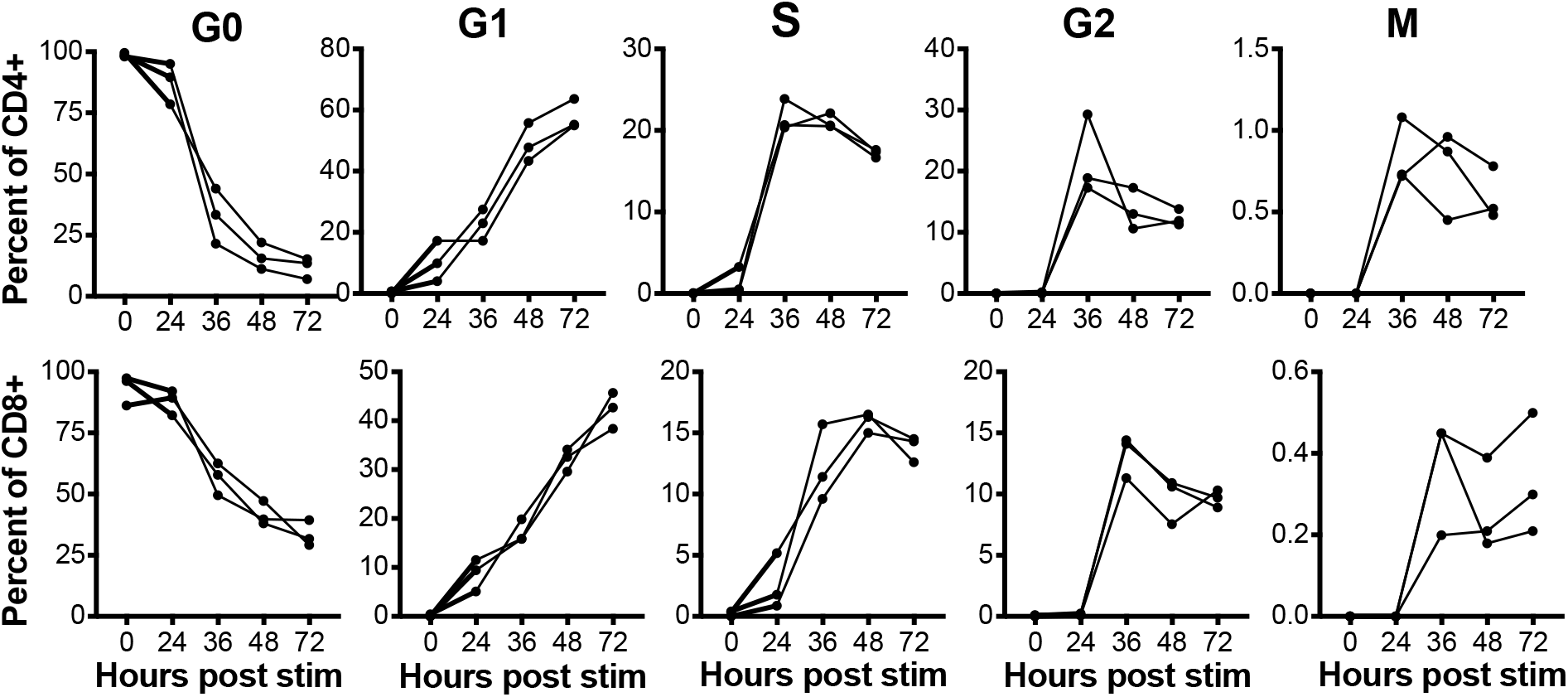
T cells enter the cell cycle following activation. Percent of CD4 (*upper panels*) or CD8 (*lower panels*) T cells in the G0, G1, S, G2, or M phases of the cell cycle as measured by Ki67 and 7AAD staining.

**Supplementary Figure 2.**
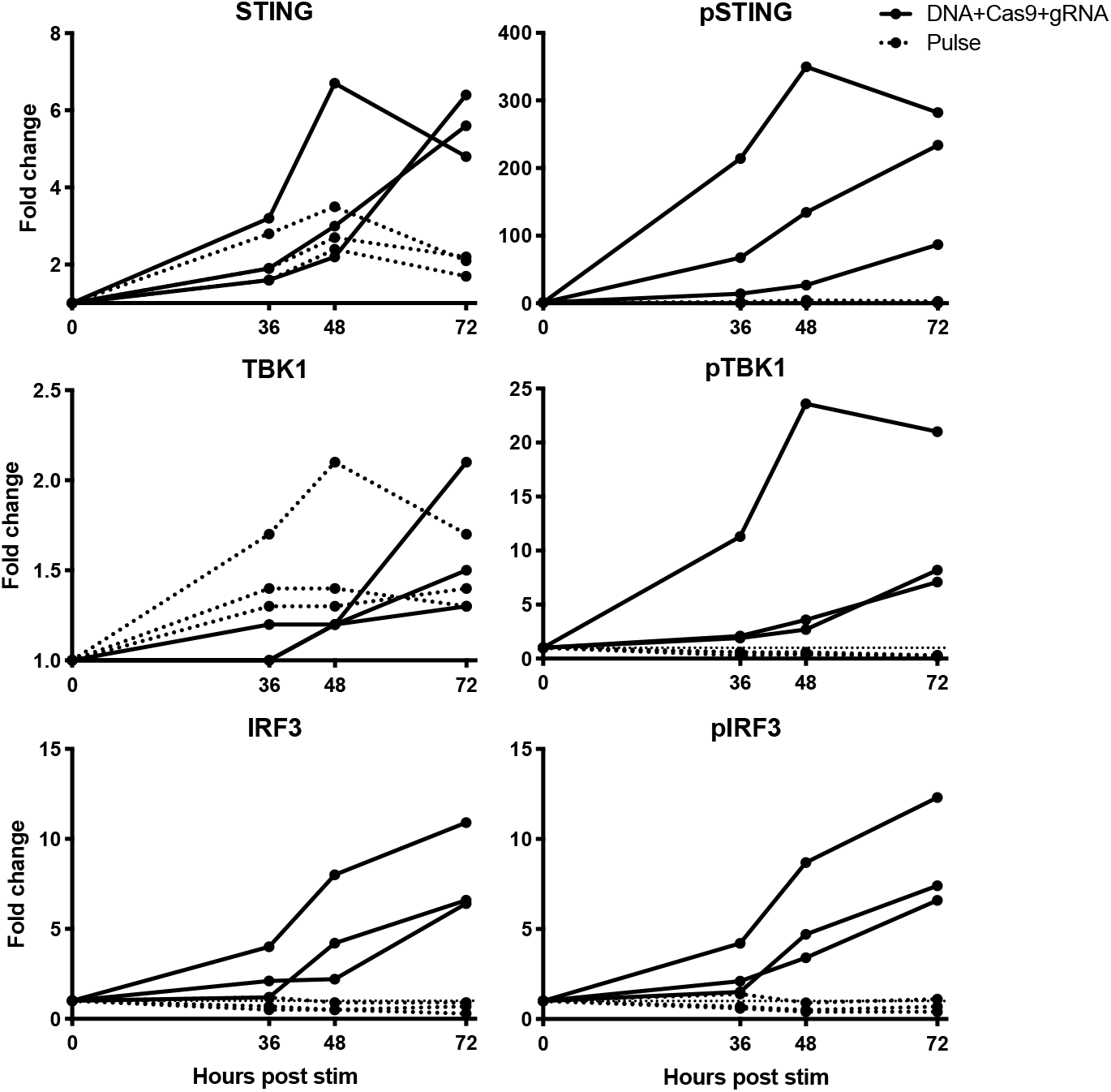
Expression of cytosolic DNA sensors increases post activation. Expression of total (*left panels*) or phosphorylated (*right panels*) STING, TBK1, and IRF3 cytosolic DNA sensors in genome engineered (*solid lines*) or control (*dashed lines*) T cells following activation as measured by Western Blot.

**Supplementary Figure 3.**
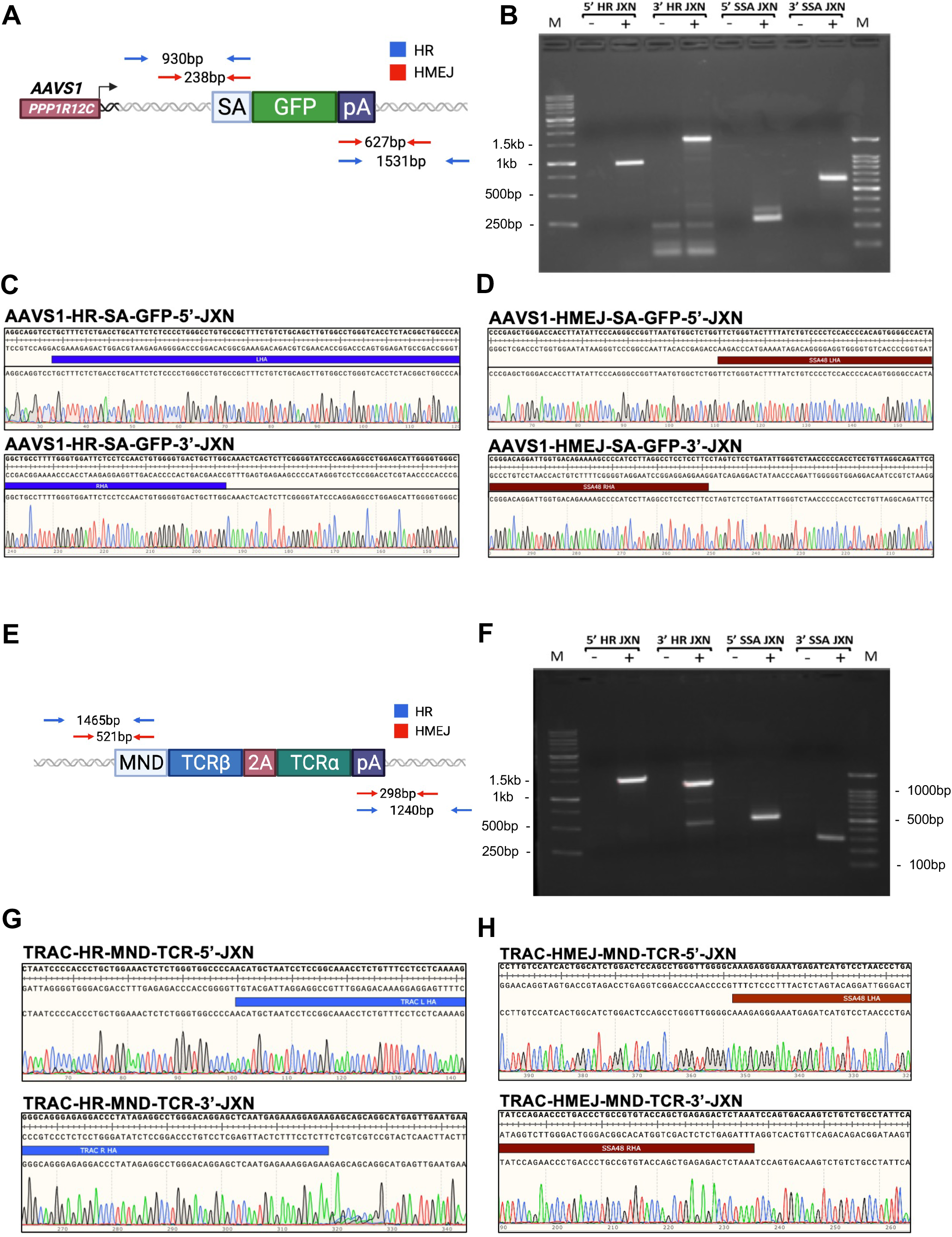
High integration fidelity at cargo-genome junction following HMEJ engineering. **(A)** Predicted sequence following integration. **(B)** Gel electrophoresis of the resultant products following PCR at the 5’ (left) and 3’ (right) junctions of the SA-GFP HR and HMEJ integration events. Gel bands were purified and samples underwent sanger sequencing. **(C,D)** Alignment of the 5’ and 3’ sanger sequencing chromatograms to the expected reference sequence. **(D,E,F,G)** Identical analysis was performed for the MND-TCR HR and HMEJ integration events.

**Supplementary Figure 4.**
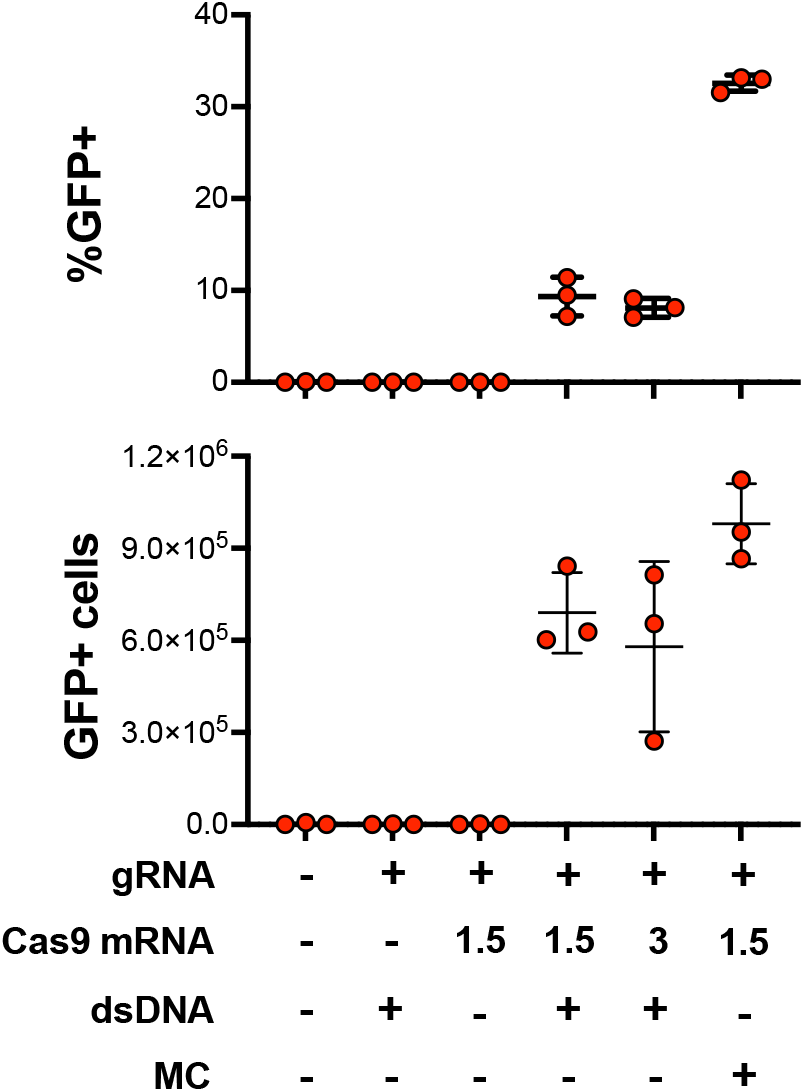
HMEJ template allows for greater integration and post-engineering viability than linear PCR template. Percentage of cells expressing GFP (*top panel*) and total number of GFP+ cells (*bottom panel*) following engineering using HMEJ or linear PCR template encoding GFP in the presence or absence of Cas9 mRNA and gRNA targeting AAVS1.

**Supplementary Figure 5.**
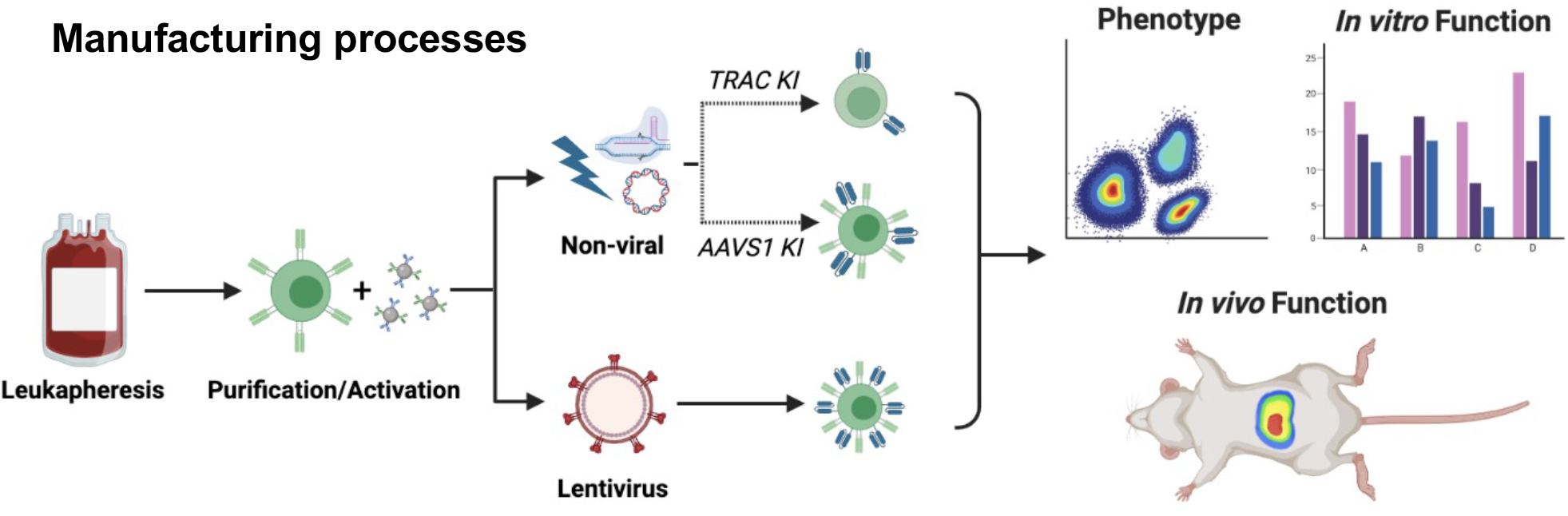
Diagram of manufacturing process for lentiviral transduced or HMEJ engineered T cells.

**Supplementary Figure 6.**
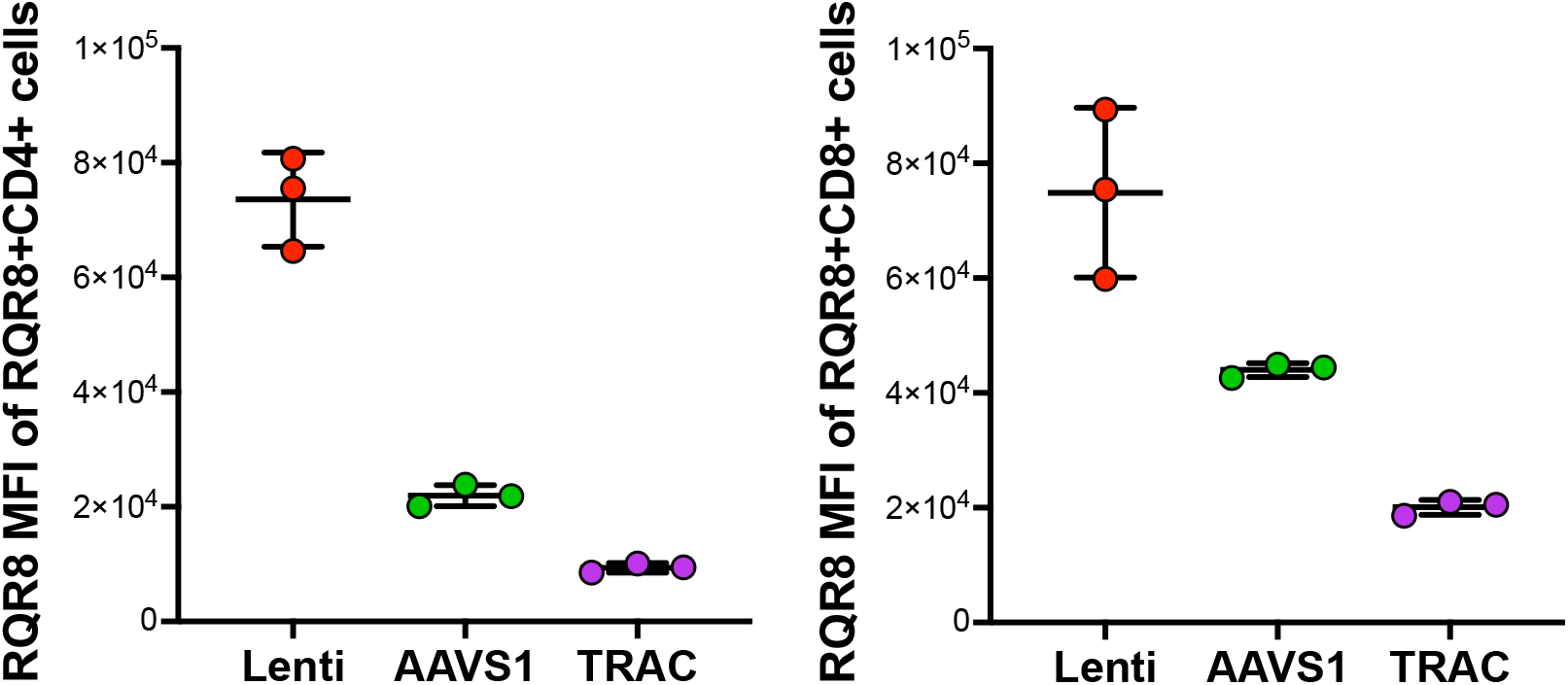
Lentiviral transduced cells have higher expression levels of CD19-CAR than non-viral engineered cells. MFI of samples following HMEJ non-viral genome engineering or lentiviral transduction with a construct encoding CD19CAR-2A-RQR8.

**Supplementary Figure 7.**
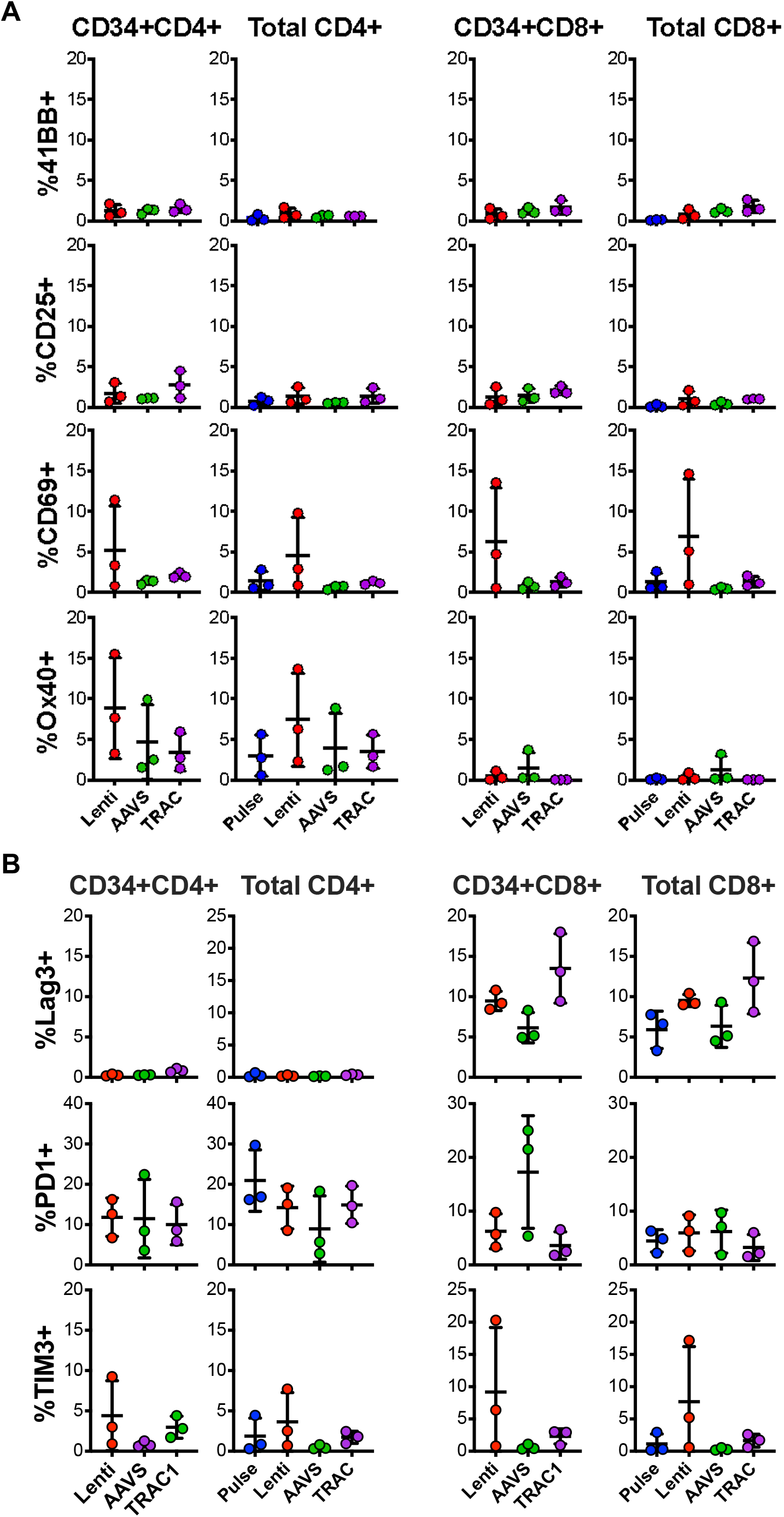
HMEJ engineered CD19-CAR T cells express similar levels of activation and exhaustion surface markers as lentiviral transduced cells. **(A)** Expression of 41BB, CD25, CD69, and Ox40 in RQR8+CD4+, total CD4+, RQR8+CD8+ and total CD8+ T cell subsets following HMEJ non-viral genome engineering or lentiviral transduction with a construct encoding CD19CAR-2A-RQR8. **(B)** Expression of Lag3, PD1, and TIM3 in RQR8+CD4+, total CD4+, RQR8+CD8+ and total CD8+ T cell subsets following HMEJ non-viral genome engineering or lentiviral transduction with a construct encoding CD19CAR-2A-RQR8. Lack of statistically significant differences was determined by comparing TRAC, AAVS1, and pulse-only cells to Lenti cells using One-way ANOVA followed by Dunnett’s multiple comparison test.

**Supplementary Figure 8.**
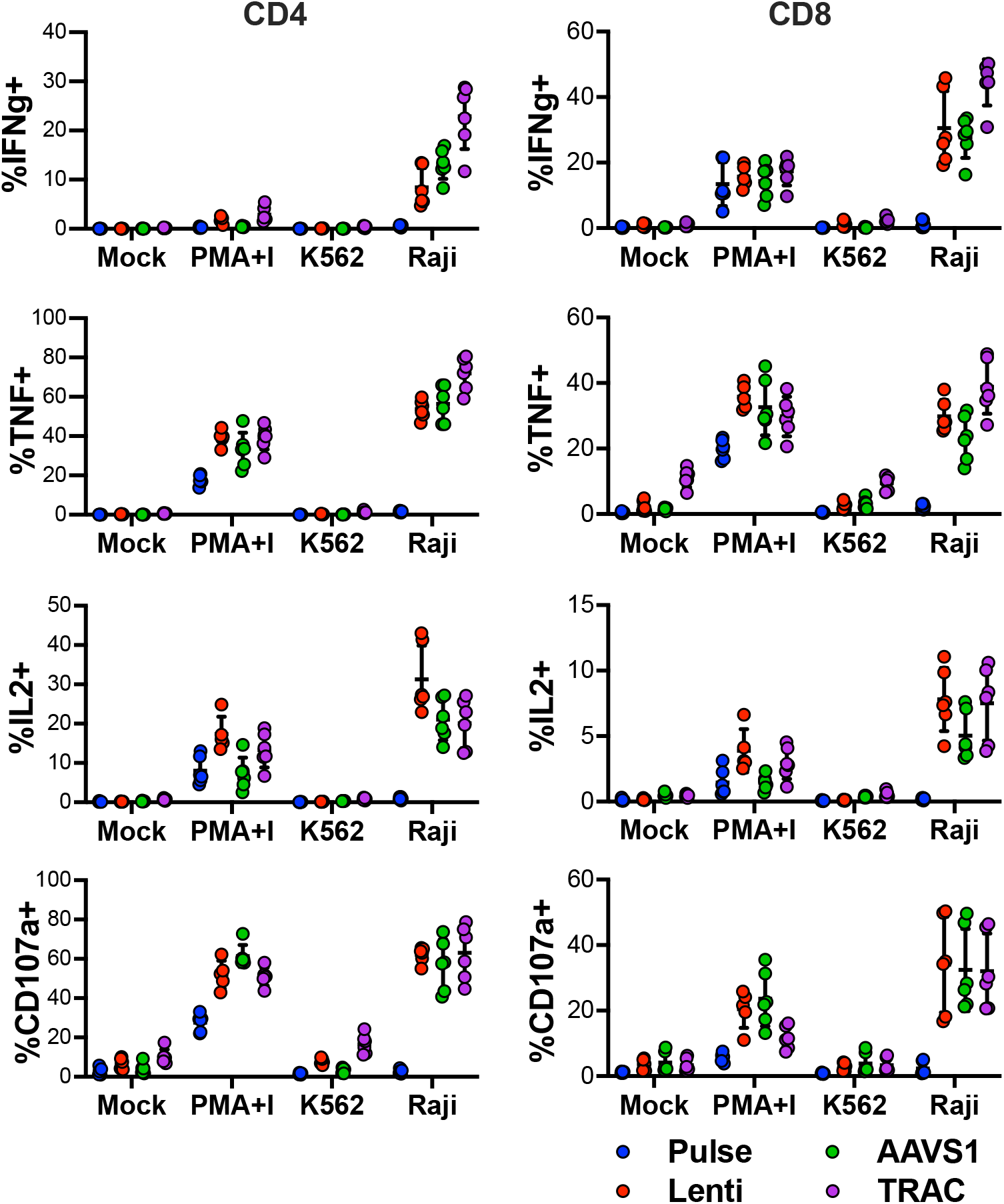
HMEJ engineered and lentiviral transduced cells CD19-CAR T cells produce cytokine in response target cells as measured by ICS. Percentage of cells expressing cytokines IFNγ, TNF, and IL2 as well as a degranulation marker CD107a in CD4 (*left panels*) and CD8 (*right panels*) CD19-CAR T cells following coculture with CD19+ Raji target cells.

**Supplementary Figure 9.**
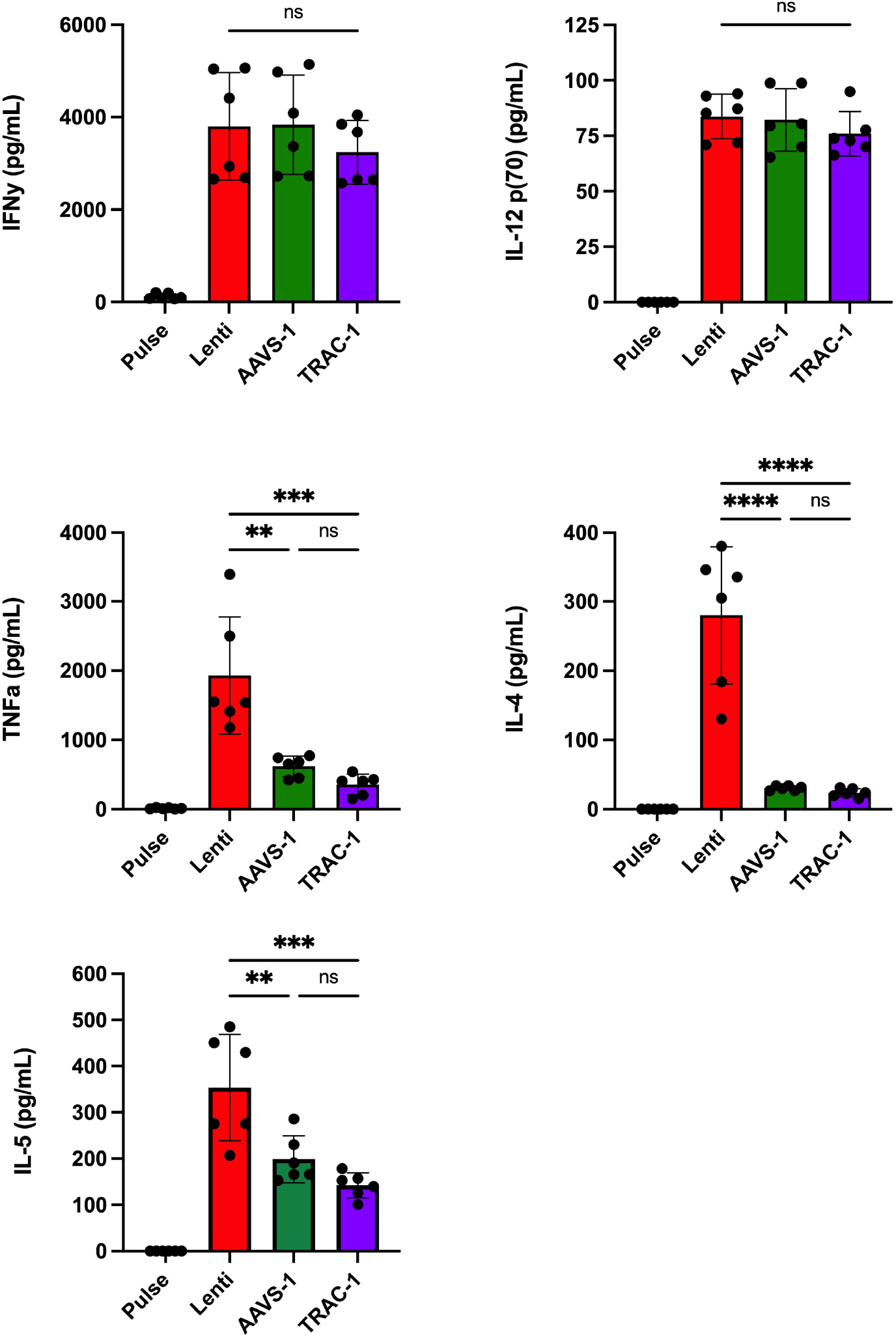
HMEJ engineered and lentiviral transduced cells CD19-CAR T cells produce cytokine in response target cells as measured by Luminex. Concentration of IFNγ, TNF, IL4, and IL5 in the supernatant of CD19-CAR T cells following coculture with CD19+ Raji target cells. All statistical analyses were done using One-way ANOVA followed by Tukey’s multiple comparison test. (n = 6 independent biological donors) (*p<0.05, **p<0.01, ***p<0.001, ****p<0.0001).

**Supplementary Table 1.**
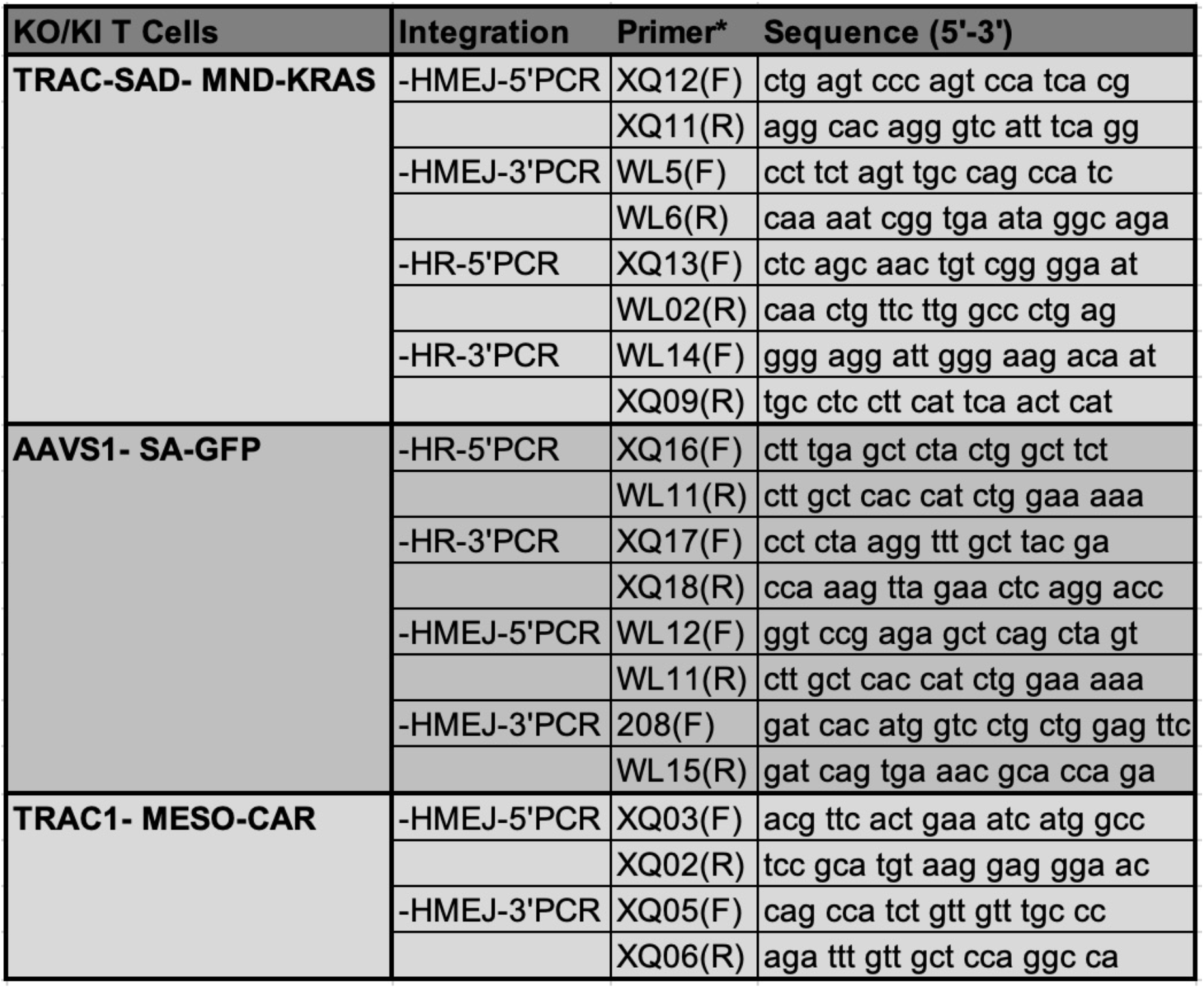
List of primers used for PCR experiments.

**Supplementary Table 2.** List of antibodies used for flow cytometry experiments.

| **Marker** | **Fluorophore** | **Clone** | **Product number** | **Company** |
| --- | --- | --- | --- | --- |
| CD3 | Cy55PE | SK7 | 35-0036-42 | Invitrogen |
| CD4 | BV605 | RPA-T4 | 300556 | BioLegend |
| CD8 | BV650 | RPA-T8 | 301042 | BioLegend |
| CD45ro | Cy7PE | UCHL1 | 304230 | BioLegend |
| CD62L | AF647 | DREG-56 | 304818 | BioLegend |
| CD107a | FITC | H4A3 | 555800 | BD Pharmingen |
| LAG3 | APC | 7H2C65 | 369212 | BioLegend |
| TIM3 | PECy7 | F38-2E2 | 345014 | BioLegend |
| PD1 | BV421 | EH12.1 | 562516 | BD Horizon |
| CD25 | BV650 | BC96 | 302634 | BioLegend |
| CD69 | PECy7 | FN50 | 310912 | BioLegend |
| Ox40 | FITC | Ber-ACT35 | 350006 | BioLegend |
| 4-1BB | PE | 4B4-1 | 309804 | BioLegend |
| Ki67 | APC | B56 | 558615 | BD Pharmingen |
| CD34 | PE | QBEND/10 | MA1-10205 | Invitrogen |
| mTCRb | Cy7PE | H57-597 | 25-5961-82 | Invitrogen |
| IFNg | eFluor450 | 4S.B3 | 48-7319-42 | Invitrogen |
| IL2 | PECy7 | MQ1-17H12 | 25-7029-42 | Invitrogen |
| TNF | APC | MAb11 | 17-7349-82 | Invitrogen |

## References

1. Khan, S. et al. Role of Recombinant DNA Technology to Improve Life. Int. J. Genomics Proteomics 2016, 2405954 (2016).

2. Spolski, R., Li, P. & Leonard, W. J. Biology and regulation of IL-2: from molecular mechanisms to human therapy. Nat. Rev. Immunol. 18, 648–659 (2018).

3. Maetzig, T., Galla, M., Baum, C. & Schambach, A. Gammaretroviral vectors: biology, technology and application. Viruses 3, 677–713 (2011).

4. Dotti, G., Gottschalk, S., Savoldo, B. & Brenner, M. K. Design and development of therapies using chimeric antigen receptor-expressing T cells. Immunol. Rev. 257, 107–126 (2014).

5. Walther, W. & Stein, U. Viral vectors for gene transfer: a review of their use in the treatment of human diseases. Drugs 60, 249–271 (2000).

6. Bulcha, J. T., Wang, Y., Ma, H., Tai, P. W. L. & Gao, G. Viral vector platforms within the gene therapy landscape. Signal Transduct Target Ther 6, 53 (2021).

7. Soundara Rajan, T., Gugliandolo, A., Bramanti, P. & Mazzon, E. In Vitro-Transcribed mRNA Chimeric Antigen Receptor T Cell (IVT mRNA CAR T) Therapy in Hematologic and Solid Tumor Management: A Preclinical Update. Int. J. Mol. Sci. 21, (2020).

8. Kebriaei, P. et al. Phase I trials using Sleeping Beauty to generate CD19-specific CAR T cells. J. Clin. Invest. 126, 3363–3376 (2016).

9. Monjezi, R. et al. Enhanced CAR T-cell engineering using non-viral Sleeping Beauty transposition from minicircle vectors. Leukemia 31, 186–194 (2017).

10. Li, X. et al. A resurrected mammalian hAT transposable element and a closely related insect element are highly active in human cell culture. Proc. Natl. Acad. Sci. U. S. A. 110, E478–87 (2013).

11. Bishop, D. C. et al. PiggyBac-Engineered T Cells Expressing CD19-Specific CARs that Lack IgG1 Fc Spacers Have Potent Activity against B-ALL Xenografts. Mol. Ther. 26, 1883–1895 (2018).

12. Nguyen, D. N. et al. Polymer-stabilized Cas9 nanoparticles and modified repair templates increase genome editing efficiency. Nat. Biotechnol. 38, 44–49 (2020).

13. Schober, K. et al. Orthotopic replacement of T-cell receptor α- and β-chains with preservation of near-physiological T-cell function. Nat Biomed Eng 3, 974–984 (2019).

14. Roth, T. L. et al. Reprogramming human T cell function and specificity with non-viral genome targeting. Nature 559, 405–409 (2018).

15. Capecchi, M. R. The new mouse genetics: altering the genome by gene targeting. Trends Genet. 5, 70–76 (1989).

16. Puchta, H., Dujon, B. & Hohn, B. Homologous recombination in plant cells is enhanced by in vivo induction of double strand breaks into DNA by a site-specific endonuclease. Nucleic Acids Res. 21, 5034–5040 (1993).

17. Anzalone, A. V., Koblan, L. W. & Liu, D. R. Genome editing with CRISPR-Cas nucleases, base editors, transposases and prime editors. Nat. Biotechnol. 38, 824–844 (2020).

18. Lee, J., Chung, J.-H., Kim, H. M., Kim, D.-W. & Kim, H. Designed nucleases for targeted genome editing. Plant Biotechnol. J. 14, 448–462 (2016).

19. Hendel, A. et al. Chemically modified guide RNAs enhance CRISPR-Cas genome editing in human primary cells. Nat. Biotechnol. 33, 985–989 (2015).

20. Webber, B. R. et al. Highly efficient multiplex human T cell engineering without double-strand breaks using Cas9 base editors. Nat. Commun. 10, 5222 (2019).

21. Johnson, M. J., Laoharawee, K., Lahr, W. S., Webber, B. R. & Moriarity, B. S. Engineering of Primary Human B cells with CRISPR/Cas9 Targeted Nuclease. Sci. Rep. 8, 12144 (2018).

22. Osborn, M. J. et al. Evaluation of TCR Gene Editing Achieved by TALENs, CRISPR/Cas9, and megaTAL Nucleases. Mol. Ther. 24, 570–581 (2016).

23. Zeng, J. et al. Therapeutic base editing of human hematopoietic stem cells. Nat. Med. 26, 535–541 (2020).

24. Bak, R. O., Dever, D. P. & Porteus, M. H. CRISPR/Cas9 genome editing in human hematopoietic stem cells. Nat. Protoc. 13, 358–376 (2018).

25. June, C. H., Blazar, B. R. & Riley, J. L. Engineering lymphocyte subsets: tools, trials and tribulations. Nat. Rev. Immunol. 9, 704–716 (2009).

26. Sather, B. D. et al. Efficient modification of CCR5 in primary human hematopoietic cells using a megaTAL nuclease and AAV donor template. Sci. Transl. Med. 7, 307ra156 (2015).

27. Osborn, M. J. et al. CRISPR/Cas9-Based Cellular Engineering for Targeted Gene Overexpression. Int. J. Mol. Sci. 19, (2018).

28. Pomeroy, E. J. et al. A Genetically Engineered Primary Human Natural Killer Cell Platform for Cancer Immunotherapy. Mol. Ther. 28, 52–63 (2020).

29. Wang, D., Tai, P. W. L. & Gao, G. Adeno-associated virus vector as a platform for gene therapy delivery. Nat. Rev. Drug Discov. 18, 358–378 (2019).

30. Samulski, R. J. & Muzyczka, N. AAV-Mediated Gene Therapy for Research and Therapeutic Purposes. Annu Rev Virol 1, 427–451 (2014).

31. Deyle, D. R., Li, L. B., Ren, G. & Russell, D. W. The effects of polymorphisms on human gene targeting. Nucleic Acids Res. 42, 3119–3124 (2014).

32. Rosenberg, S. A. & Restifo, N. P. Adoptive cell transfer as personalized immunotherapy for human cancer. Science 348, 62–68 (2015).

33. Chandran, S. S. & Klebanoff, C. A. T cell receptor-based cancer immunotherapy: Emerging efficacy and pathways of resistance. Immunol. Rev. 290, 127–147 (2019).

34. Semenova, N. et al. Multiple cytosolic DNA sensors bind plasmid DNA after transfection. Nucleic Acids Res. 47, 10235–10246 (2019).

35. Maurisse, R. et al. Comparative transfection of DNA into primary and transformed mammalian cells from different lineages. BMC Biotechnol. 10, 9 (2010).

36. Chen, Q., Sun, L. & Chen, Z. J. Regulation and function of the cGAS-STING pathway of cytosolic DNA sensing. Nat. Immunol. 17, 1142–1149 (2016).

37. Paludan, S. R. & Bowie, A. G. Immune sensing of DNA. Immunity 38, 870–880 (2013).

38. Wu, J. & Chen, Z. J. Innate immune sensing and signaling of cytosolic nucleic acids. Annu. Rev. Immunol. 32, 461–488 (2014).

39. Clark, K., Plater, L., Peggie, M. & Cohen, P. Use of the pharmacological inhibitor BX795 to study the regulation and physiological roles of TBK1 and IkappaB kinase epsilon: a distinct upstream kinase mediates Ser-172 phosphorylation and activation. J. Biol. Chem. 284, 14136–14146 (2009).

40. Richters, A. et al. Identification and further development of potent TBK1 inhibitors. ACS Chem. Biol. 10, 289–298 (2015).

41. Decout, A., Katz, J. D., Venkatraman, S. & Ablasser, A. The cGAS-STING pathway as a therapeutic target in inflammatory diseases. Nat. Rev. Immunol. 21, 548–569 (2021).

42. Tsukahara, T. et al. CD19 target-engineered T-cells accumulate at tumor lesions in human B-cell lymphoma xenograft mouse models. Biochem. Biophys. Res. Commun. 438, 84–89 (2013).

43. Hu, S.-I. et al. Pre-clinical assessment of chimeric antigen receptor t cell therapy targeting CD19+ B cell malignancy. Ann Transl Med 8, 584 (2020).

44. Zhang, Z., Qiu, S., Zhang, X. & Chen, W. Optimized DNA electroporation for primary human T cell engineering. BMC Biotechnol. 18, 4 (2018).

45. Kay, M. A., He, C.-Y. & Chen, Z.-Y. A robust system for production of minicircle DNA vectors. Nat. Biotechnol. 28, 1287–1289 (2010).

46. Obst, R. The Timing of T Cell Priming and Cycling. Front. Immunol. 6, 563 (2015).

47. Yu, L. & Liu, P. Cytosolic DNA sensing by cGAS: regulation, function, and human diseases. Signal Transduct Target Ther 6, 170 (2021).

48. Vance, R. E. Cytosolic DNA Sensing: The Field Narrows. Immunity vol. 45 227–228 (2016).

49. Zahid, A., Ismail, H., Li, B. & Jin, T. Molecular and Structural Basis of DNA Sensors in Antiviral Innate Immunity. Front. Immunol. 11, 613039 (2020).

50. Wierson, W. A. et al. Efficient targeted integration directed by short homology in zebrafish and mammalian cells. Elife 9, (2020).

51. Xue, C. & Greene, E. C. DNA Repair Pathway Choices in CRISPR-Cas9-Mediated Genome Editing. Trends Genet. 37, 639–656 (2021).

52. Li, X. et al. Efficient SSA-mediated precise genome editing using CRISPR/Cas9. FEBS J. 285, 3362–3375 (2018).

53. Bewg, W. P., Ci, D. & Tsai, C.-J. Genome Editing in Trees: From Multiple Repair Pathways to Long-Term Stability. Front. Plant Sci. 9, 1732 (2018).

54. Yao, X. et al. Homology-mediated end joining-based targeted integration using CRISPR/Cas9. Cell Res. 27, 801–814 (2017).

55. Dahlman, J. E. et al. Orthogonal gene knockout and activation with a catalytically active Cas9 nuclease. Nat. Biotechnol. 33, 1159–1161 (2015).

56. Philip, B. et al. A highly compact epitope-based marker/suicide gene for easier and safer T-cell therapy. Blood 124, 1277–1287 (2014).

57. Berger, A. Th1 and Th2 responses: what are they? BMJ 321, 424 (2000).

58. Eyquem, J. et al. Targeting a CAR to the TRAC locus with CRISPR/Cas9 enhances tumour rejection. Nature 543, 113–117 (2017).

59. DeRenzo, C. & Gottschalk, S. Genetic Modification Strategies to Enhance CAR T Cell Persistence for Patients With Solid Tumors. Front. Immunol. 10, 218 (2019).

60. Liu, X. et al. A Chimeric Switch-Receptor Targeting PD1 Augments the Efficacy of Second-Generation CAR T Cells in Advanced Solid Tumors. Cancer Res. 76, 1578–1590 (2016).

61. Yeku, O. O., Purdon, T. J., Koneru, M., Spriggs, D. & Brentjens, R. J. Armored CAR T cells enhance antitumor efficacy and overcome the tumor microenvironment. Sci. Rep. 7, 10541 (2017).

62. Yu, S., Yi, M., Qin, S. & Wu, K. Next generation chimeric antigen receptor T cells: safety strategies to overcome toxicity. Mol. Cancer 18, 125 (2019).

63. Brandt, L. J. B., Barnkob, M. B., Michaels, Y. S., Heiselberg, J. & Barington, T. Emerging Approaches for Regulation and Control of CAR T Cells: A Mini Review. Front. Immunol. 11, 326 (2020).

64. Srivastava, S. et al. Logic-Gated ROR1 Chimeric Antigen Receptor Expression Rescues T Cell-Mediated Toxicity to Normal Tissues and Enables Selective Tumor Targeting. Cancer Cell 35, 489–503.e8 (2019).

65. Raes, L., De Smedt, S. C., Raemdonck, K. & Braeckmans, K. Non-viral transfection technologies for next-generation therapeutic T cell engineering. Biotechnol. Adv. 49, 107760 (2021).

66. Odé, Z., Condori, J., Peterson, N., Zhou, S. & Krenciute, G. CRISPR-Mediated Non-Viral Site-Specific Gene Integration and Expression in T Cells: Protocol and Application for T-Cell Therapy. Cancers 12, (2020).

67. Pearce, E. L., Poffenberger, M. C., Chang, C.-H. & Jones, R. G. Fueling immunity: insights into metabolism and lymphocyte function. Science 342, 1242454 (2013).

68. Paludan, S. R., Reinert, L. S. & Hornung, V. DNA-stimulated cell death: implications for host defence, inflammatory diseases and cancer. Nat. Rev. Immunol. 19, 141–153 (2019).

69. Ceccaldi, R., Rondinelli, B. & D’Andrea, A. D. Repair Pathway Choices and Consequences at the Double-Strand Break. Trends Cell Biol. 26, 52–64 (2016).

70. Schweizer, M. & Merten, O.-W. Large-scale production means for the manufacturing of lentiviral vectors. Curr. Gene Ther. 10, 474–486 (2010).

71. Merten, O.-W., Hebben, M. & Bovolenta, C. Production of lentiviral vectors. Mol Ther Methods Clin Dev 3, 16017 (2016).

72. Alves, C. P. A., Prazeres, D. M. F. & Monteiro, G. A. Minicircle biopharmaceuticals–an overview of purification strategies. Front. Chem. Eng. Chin. 2, (2021).

73. Tian, Y., Li, Y., Shao, Y. & Zhang, Y. Gene modification strategies for next-generation CAR T cells against solid cancers. J. Hematol. Oncol. 13, 54 (2020).

74. Nakamura, K. et al. Autologous antigen-presenting cells efficiently expand piggyBac transposon CAR-T cells with predominant memory phenotype. Mol Ther Methods Clin Dev 21, 315–324 (2021).

75. 75. Unforeseen Development of Lymphoma Derived from piggyBac CAR T Cells. Cancer Discov. 11, 1613 (2021).

76. Lukjanov, V., Koutná, I. & Šimara, P. CAR T-Cell Production Using Nonviral Approaches. J Immunol Res 2021, 6644685 (2021).

77. Chicaybam, L. et al. Transposon-mediated generation of CAR-T cells shows efficient anti B-cell leukemia response after ex vivo expansion. Gene Ther. 27, 85–95 (2020).

78. Kaiser, A. D. et al. Towards a commercial process for the manufacture of genetically modified T cells for therapy. Cancer Gene Ther. 22, 72–78 (2015).

79. Wang, X. & Rivière, I. Clinical manufacturing of CAR T cells: foundation of a promising therapy. Mol Ther Oncolytics 3, 16015 (2016).

80. Gee, A. P. GMP CAR-T cell production. Best Pract. Res. Clin. Haematol. 31, 126–134 (2018).

